# PRRT2 as an auxiliary regulator of Nav channel slow inactivation

**DOI:** 10.1101/2025.09.27.678785

**Authors:** Bin Lu, Qi-Wu Xu, Jing Zhang, Xue-Mei Wu, Jun-Yan He, Jing-Qiu Peng, Guang Yang, Ke-Xian Li, Ling Zhuang, Yu-Xian Zhang, Zhi-Ying Wu, Zhi-Qi Xiong

## Abstract

During sustained activity, voltage-gated sodium (Nav) channels enter a slow-inactivated state to limit cellular hyperexcitability. Disruption of this regulatory process has been implicated in skeletal, cardiac and neurological disorders. While the kinetics of this process are well characterized, its endogenous modulators remain unclear. Here, we identify Proline-Rich Transmembrane Protein 2 (PRRT2) as a native regulator of Nav channel slow inactivation. We show that PRRT2 facilitates the entry of Nav channels into the slow-inactivated state and delays their recovery, a regulatory effect conserved from zebrafish to humans. PRRT2 forms molecular complexes with Nav channels both *in vitro* and *in vivo*. In the mouse cortex, PRRT2 deficiency impairs the slow inactivation of Nav channels in neuronal axons, leading to reduced cortical resilience in response to hyperexcitable challenges. Together, these findings establish PRRT2 as a physiological modulator of Nav channel slow inactivation and reveal a mechanism that supports cortical resilience to pathological perturbations.

## Introduction

Voltage-gated sodium (Nav) channels are critical for the initiation and propagation of electrical signals in excitable cells, contributing to cellular excitability in various tissues(Hodgkin & Huxley, 1952; Ulbricht, 1977). In mammals, including humans, Nav channels consist of nine α-subunit isoforms (Nav1.1-Nav1.9), which are differentially expressed and distributed across different tissues to regulate specific physiological functions(Catterall, Goldin, & Waxman, 2005). Mutations in Nav channels that lead to either loss- or gain-of-function of these channels are associated with a wide spectrum of diseases, including neurological disorders(Escayg & Goldin, 2010; Hedrich, Lauxmann, & Lerche, 2019; Meisler, Hill, & Yu, 2021), pain syndromes(Dib-Hajj, Geha, & Waxman, 2017; Waxman, 2013), muscle illnesses(Cannon, 2018; Mantegazza, Cestele, & Catterall, 2021) and cardiac diseases(Remme, 2013). Consequently, the Nav channels are considered promising therapeutic targets for treating these conditions(Noreng, Li, & Payandeh, 2021; Wisedchaisri & Gamal El-Din, 2022).

Voltage-dependent conformational changes enable Nav channels to transition between resting, open and inactivated states(Catterall, Wisedchaisri, & Zheng, 2020). Eukaryotic Nav channels undergo two major forms of inactivation (fast and slow inactivation), which are distinguished by their kinetics of onset and recovery(Goldin, 2003). Fast inactivation is triggered within milliseconds by brief depolarization and is rapidly reversed upon hyperpolarization, typically within tens of milliseconds (Armstrong & Bezanilla, 1977; Bezanilla & Armstrong, 1977). In contrast, slow inactivation develops during prolonged depolarizations and the slow inactivated channels require substantially longer periods to recover, ranging from hundreds of milliseconds to minutes(Rudy, 1978). Previous studies have shown that slow inactivation regulates Nav channel availability during sustained depolarization or high-frequency repetitive activity, thereby modulating the cellular excitability(Goldin, 2003; Silva, 2014; Vilin & Ruben, 2001). Impairment of this regulatory process due to mutations in Nav channels has been associated with disorders such as periodic paralysis(Hayward, Sandoval, & Cannon, 1999), cardiac arrhythmias(Richmond, Featherstone, Hartmann, & Ruben, 1998; Vilin, Makita, George, & Ruben, 1999) and epilepsy(Ghovanloo et al., 2023).

Beyond the intrinsic hierarchical gating mechanisms, auxiliary regulators further modulate Nav channel inactivation dynamics(Goldin, 2003). For example, β subunits of sodium channels (SCN1B) regulate inactivation of Nav channels in the heterologous expression system, although their effects vary across different Nav isoforms and cell types(He & Soderlund, 2014; Vilin et al., 1999; Webb, Wu, & Cannon, 2009). Fibroblast growth factor homologous factors (FHFs) interact with the carboxy terminus of Nav channels(Gade, Marx, & Pitt, 2020; Goetz et al., 2009), promoting intermediated or long-term inactivation, with recovery times spanning hundreds of milliseconds(Dover, Solinas, D’Angelo, & Goldfarb, 2010; Venkatesan, Liu, & Goldfarb, 2014). Although these auxiliary proteins have been identified as regulators of fast or intermediate inactivation, native modulators that specifically target slow inactivation of Nav channels remain largely unknown.

Recent studies on proline-rich transmembrane protein 2 (PRRT2) provide new insights into this issue. The *PRRT2*, a member of the Dispanins B subfamily (DspB)(Sallman Almen, Bringeland, Fredriksson, & Schioth, 2012), has been identified as a causative gene for paroxysmal kinesigenic dyskinesia(W. J. Chen et al., 2011; Lee et al., 2012; J. L. Wang et al., 2011). *In vitro* studies have shown that PRRT2 regulates Nav channel surface expression and delays recovery of these channels from inactivated states induced by brief depolarization(Fruscione et al., 2018; Lu et al., 2021; Valente et al., 2023). However, these studies did not clearly distinguish between the effects of PRRT2 on fast versus slow inactivation, nor did they provide direct *in vivo* evidence for its role in regulating Nav channel inactivation. Our original observations that PRRT2 expression increased the accumulation of inactivated Nav channels during repetitive stimulation(Lu et al., 2021) suggest that PRRT2 may be a previously unrecognized auxiliary regulator of Nav channel slow inactivation.

In this study, we combined whole-cell voltage-clamp recordings, co-immunoprecipitation, and *in vivo* approaches to investigate the role of PRRT2 in regulating Nav channel slow inactivation, and to examine how deficits in PRRT2-mediated modulation of this process affects neuronal function in awake mice.

## Results

### PRRT2 regulates the slow inactivation of Nav1.2 channels

Fast and slow inactivation of Nav channels proceed through distinct gating pathways(Goldin, 2003). Although previous studies have implicated PRRT2 in the regulation of Nav channel inactivation(Fruscione et al., 2018; Lu et al., 2021), it remains unclear whether PRRT2 primarily influences fast inactivation, slow inactivation or both. To address this, we expressed mouse PRRT2 in HEK293T cells stably expressing Nav1.2 (Figure 1A) and applied classic electrophysiological paradigms to distinguish between the two inactivation processes.

**Figure 1.**
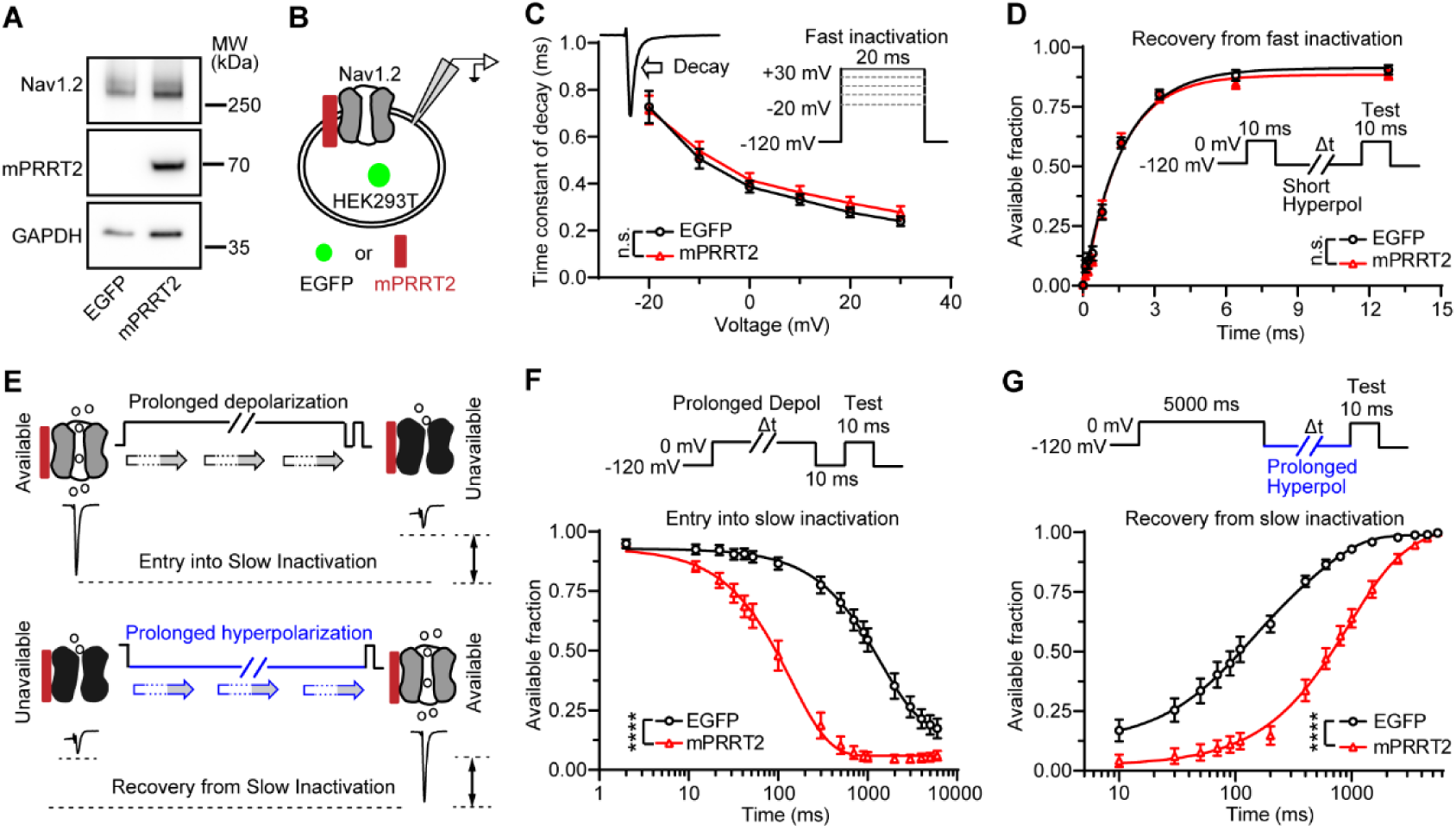
PRRT2 regulates slow inactivation of Nav1.2 channels. **(A and B)** Representative immunoblots showing PRRT2 protein expression (A) and schematic of whole-cell recording (B) in Nav1.2-stably expressing HEK293T cells transfected with constructs encoding either EGFP or mouse PRRT2 (mPRRT2). MW, molecular weight. **(C)** Fast-inactivation decay time constants of sodium currents evoked by 20-ms depolarizing steps to the indicated command voltages (EGFP: n = 13 cells, mPRRT2: n = 13 cells). Insert: representative sodium current traces elicited by a brief depolarization (left) and voltage-clamp protocol used to assess fast inactivation (right). The arrow indicates the decay of the sodium current attributable to fast inactivation. **(D)** Recovery from fast inactivation of Nav1.2 channels (EGFP: n = 10 cells, mPRRT2: n = 7 cells). Insert: Protocol for examining fast inactivation recovery. **(E)** Schematic diagram of entry into and recovery from slow inactivation of voltage-gated sodium channels. Slow inactivation modulates the availability of the sodium channels. **(F)** Entry into slow-inactivated state of Nav1.2 channels induced by progressively longer depolarization (EGFP: n = 6 cells, mPRRT2: n = 6 cells). **(G)** The effects of PRRT2 on the recovery of Nav1.2 channels from slow-inactivated state induced by 5-s depolarization (EGFP: n = 6 cells, PRRT2: n = 6 cells). Data are presented as mean ± s.e.m. Two-tailed, unpaired Student’s t-test was used in C, and two-way ANOVAs were used in D, F, and G. ****P < 0.0001. n.s., not significant.

Fast inactivation of Nav channels is rapidly triggered by brief depolarization and manifests as a current decay following a transient pore opening (Figure 1B and 1C), consistent with previous reports(Misra, Kahlig, & George, 2008; Thompson et al., 2023). To evaluate the kinetics of fast inactivation, we measured the time constant of current decay across a range of test potentials and found that PRRT2 expression did not significantly alter the rate at which Nav channels entered the fast-inactivated state (Figure 1C). Moreover, the majority of Nav1.2 channels recovered from fast inactivation within ∼10 ms at a hyperpolarized potential of -120 mV, and PRRT2 expression had a minimal effect on this recovery (Figure 1D).

In contrast to fast inactivation, Nav channel slow inactivation develops and recovers over much longer timescales (Figure 1E), typically ranging from hundreds of milliseconds to seconds, or even to minutes(Silva, 2014). Because slow-inactivated channels do not recover during brief hyperpolarization, they can be functionally distinguished from fast-inactivated channels using a classic two-pulse protocol. In this approach, a brief hyperpolarization step (e.g., -120 mV, 10 ms) is inserted between a conditioning pulse and a test pulse to recover fast-inactivated channels. Channels that have entered the slow-inactivated state remain unavailable during the test pulse, resulting in a reduction in peak sodium current (Figure 1E)(Feather-stone, Richmond, & Ruben, 1996).

To assess the development of slow inactivation, we applied depolarization pulses of variable duration, followed by a fixed 10-ms hyperpolarization before the test pulse (Figure 1F). We found that a 5-second depolarization at 0 mV was sufficient to drive most Nav1.2 channels into the slow-inactivated state (Figure 1F), consistent with previous studies(Ganguly, Thompson, & George, 2021). PRRT2 significantly promoted the entry of Nav1.2 channels into the slow-inactivated state during sustained depolarization (Figure 1F).

To examine the role of PRRT2 in channel recovery from slow inactivation, we used a protocol in which a fixed 5-second depolarization at 0 mV was followed by varying durations of hyperpolarization at -120 mV to allow channel repriming (Figure 1G). PRRT2 expression markedly slowed the recovery of Nav channels from the slow-inactivated state (Figure 1G).

To further investigate whether the regulation of PRRT2 in slow inactivation is voltage dependent, we performed a steady-state slow inactivation protocol. In this assay, 10-s depolarizing steps from -110 to -10 mV were applied to drive Nav channels into steady state, followed by a fixed hyperpolarization and a test pulse to assess channel availability (Figure 1-figure supplement 1A). We found that PRRT2 promotes the transition of Nav channels into the slow-inactivated state across a broad range of membrane potentials. PRRT2 expression induced a pronounced hyperpolarizing shift in the inactivation curve, with a V_0.5_ of -76.27 mV compared to - 48.05 mV in controls (Figure 1-figure supplement 1B).

In addition to prolonged depolarization, high-frequency repetitive depolarization can also induce Nav channel slow inactivation(Mickus, Jung, & Spruston, 1999). To determine whether PRRT2 regulates slow inactivation under these conditions, we applied a pulse-train protocol with an inter-pulse recovery interval of 10 ms. The use-dependent attenuation of peak sodium current was quantified as the fraction of current remaining across successive depolarizations (Figure 1-figure supplement 1C). Compared with the EGFP control, PRRT2 expression produced a greater decline in peak current during the train, consistent with enhanced slow inactivation during repetitive depolarizations (Figure 1-figure supplement 1C). Together, these findings indicate that PRRT2 is a potent regulator of Nav1.2 channel slow inactivation *in vitro*.

### Functional auxiliary factors in the regulation of Nav channel slow inactivation

Several auxiliary factors of Nav channels have been identified, including the SCN1B and FHF2A (also known as FGF13a), both of which are implicated in modulating Nav channel inactivation(Dover et al., 2010; Isom et al., 1992). In this study, we compared the effects of mouse SCN1B and FGF13a with those of PRRT2 on the development and recovery of Nav channel slow inactivation (Figure 2A).

**Figure 2.**
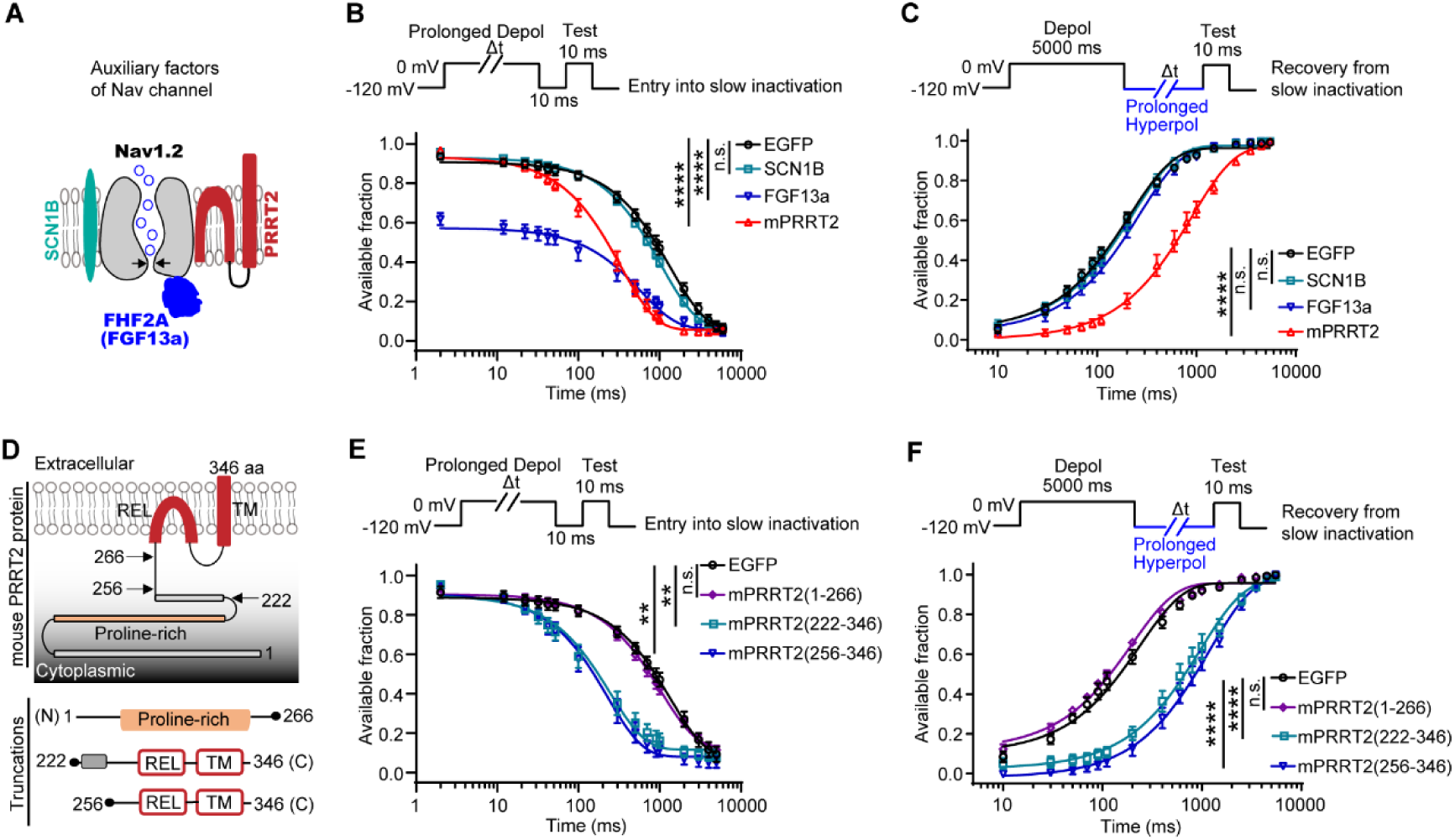
Functional auxiliary factors in the regulation of Nav channel slow inactivation. **(A)** Diagram of Nav channel with potent auxiliary factors, encompassing SCN1B, FHF2A (also known as FGF13a) and PRRT2. **(B)** The effects of auxiliary factors on the entry of Nav1.2 channels into slow inactivation (EGFP: n = 12 cells, mPRRT2: n = 14 cells, SCN1B: n = 9 cells, FGF13a: n = 12 cells). **(C)** The effects of auxiliary factors on the recovery of Nav1.2 channels from slow-inactivated state induced by 5-s depolarization (EGFP: n = 12 cells, mPRRT2: n = 12 cells, SCN1B: n = 12 cells, FGF13a: n = 12 cells). **(D)** Schematic representation of the putative topology of mouse PRRT2 and three truncations. Numbers indicate the positions of the amino acid. REL, re-entrant loop. TM, transmembrane domain. N, amino terminus. C, carboxyl terminus. aa, amino acid. **(E)** Effects of PRRT2 truncations on the entry of Nav1.2 channels into slow inactivation (EGFP: n = 5 cells, mPRRT2(1–266): n = 14 cells, mPRRT2(222–346): n = 12 cells, mPRRT2(256–346): n = 12 cells). **(F)** Effects of PRRT2 truncations on the recovery of Nav1.2 channels from slow-inactivated state induced by 5-s depolarization (EGFP: n = 5 cells, mPRRT2(1–266): n = 10 cells, mPRRT2(222–346): n = 10 cells, mPRRT2(256–346): n = 12 cells). Data are presented as mean ± s.e.m. The main effect of group was assessed using two-way ANOVAs (B, C, E and F). **P < 0.01, ****P < 0.0001, n.s., not significant.

We found that SCN1B had negligible effects on both the entry into and recovery from the slow-inactivated state of Nav1.2 compared to controls (Figures 2B and C). In contrast, FGF13a markedly accelerated the onset of inactivation (Figure 2B), consistent with previous reports(Dover et al., 2010; Venkatesan et al., 2014). Notably, FGF13a had minimal effect on the recovery of Nav channels from the slow-inactivated state induced by a 5-second depolarization (Figure 2C). These distinct effects of FGF13a on inactivation are consistent with the idea that FGF13a does not primarily modulate classical slow inactivation, but rather promotes a form of rapid-onset and long-term inactivation(Dover et al., 2010; Venkatesan et al., 2014). By comparison, PRRT2 uniquely regulated slow inactivation by both facilitating entry into and delaying recovery from the slow-inactivated state of Nav channels (Figures 2B and C). The rodent *Prrt2* gene encodes a 346-amino-acid protein containing a proline-rich domain in the N-terminus, a re-entrant loop, and a transmembrane domain in the C-terminus (Figure 2D). To identify the functional domain(s) responsible for regulating slow inactivation, we generated a series of protein-truncating variants (Figure 2D).

We first examined PRRT2(222–346), which lacks the N-terminal proline-rich domain (Figure 2D). This variant retained activity similar to the full-length PRRT2, enhancing slow inactivation development and delaying recovery (Figures 2E and F). A shorter construct, PRRT2(256–346), which retains the intact C-terminus, also preserved regulatory function in modulating slow inactivation compared to the EGFP control (Figures 2E and F). To further test whether the C-terminal region is necessary for the effect of PRRT2 on Nav channel slow inactivation, we examined PRRT2(1–266), a truncated variant lacking the membrane-associated C-terminal domains (Figure 2D). Unlike variants containing the C-terminal region, PRRT2(1–266) showed no detectable effect on either the development or recovery of Nav channel slow inactivation compared to the control (Figures 2E and F). Notably, this lack of effect was not attributable to absent expression of the truncated construct (Figure 2-figure supplement 1A and B). Together, these results indicate that the C-terminal region of PRRT2, including the re-entrant loop and transmembrane domain, is both necessary and sufficient to mediate its regulatory effect on slow inactivation of Nav1.2 channels *in vitro*.

### Evolutionarily conserved effects of PRRT2 on Nav channel slow inactivation

The *PRRT2* and *PRRT2*-like genes emerged in vertebrates and have been conserved throughout evolution(Sallman Almen et al., 2012). The alignment of amino acid sequences from zebrafish, mouse, and human PRRT2 revealed 77.46% identity between human and mouse PRRT2, and 43.60% identity between zebrafish and mouse PRRT2 (Figures 3A and B). Notably, the majority of conserved residues are located within the C-terminal region of the PRRT2 proteins (Figure 3A). Given the essential role of the C-terminus of mouse PRRT2 in regulating Nav channel slow inactivation, we examined whether PRRT2 proteins from different species exhibit similar effects.

**Figure 3.**
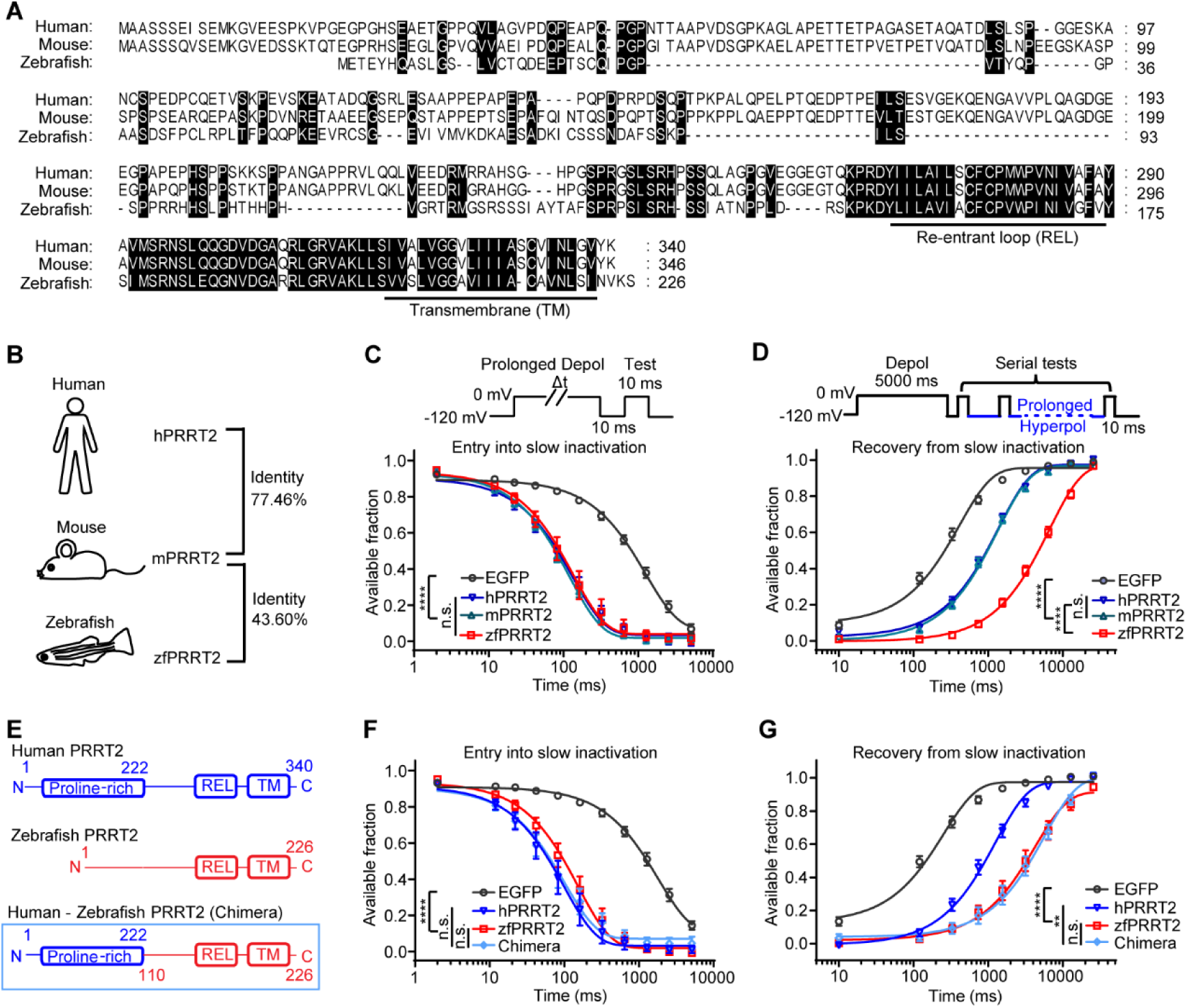
Evolutionarily conserved effects of PRRT2 on Nav channel slow inactivation. **(A)** Sequence alignment of PRRT2 protein in human, mouse and zebrafish. Conserved amino acids are highlighted and the membrane-associated domains in carboxyl terminus of PRRT2 are underlined. REL, re-entrant loop. TM, transmembrane domain. **(B)** Sequence identity of PRRT2 protein in human (hPRRT2), mouse (mPRRT2) and zebrafish (zfPRRT2). Identity is calculated by using protein BLAST tool from NCBI. **(C)** The effects of PRRT2 from different species on the entry of Nav1.2 channels into slow inactivation (EGFP: n = 16 cells, mPRRT2: n = 19 cells, hPRRT2: n = 20 cells, zfPRRT2: n = 11 cells). **(D)** The effects of PRRT2 from different species on the recovery of Nav1.2 channels from slow-inactivated state induced by 5-s depolarization (EGFP: n = 16 cells, mPRRT2: n = 19 cells, hPRRT2: n = 19 cells, zfPRRT2: n = 11 cells). **(E)** Diagram for the chimeric construct of PRRT2 originated from human and zebrafish. **(F)** The effect of chimeric PRRT2 on the entry of Nav1.2 channels into slow inactivation (EGFP: n = 11 cells, hPRRT2: n = 6 cells, zfPRRT2: n = 10 cells, Chimera: n = 11 cells). **(G)** The effect of chimeric PRRT2 on the recovery of Nav1.2 channels from slow-inactivated state induced by 5-s depolarization (EGFP: n = 11 cells, hPRRT2: n = 6 cells, zfPRRT2: n = 10 cells, Chimera: n = 9 cells). Data are presented as mean ± s.e.m. The main effect of group was assessed using two-way ANOVAs (C, D, F and G). **P < 0.01, ****P < 0.0001, n.s., not significant.

By comparing the effects of zebrafish, mouse, and human PRRT2 on Nav channel slow inactivation, we found that all three PRRT2 orthologs similarly promoted the entry of Nav1.2 channels into the slow-inactivated state (Figure 3C). Notably, in recovery assays, zebrafish PRRT2 was more effective at delaying the recovery process compared to its mammalian counterparts (Figure 3D). To identify the domain responsible for this enhanced effect, we generated a chimeric PRRT2 construct by fusing the C-terminal region of zebrafish PRRT2 to the N-terminal region of human PRRT2 (Figure 3E). The chimeric PRRT2 exhibited comparable effects on the rate of slow inactivation development to those observed with both human and zebrafish PRRT2 (Figure 3F). Notably, the chimeric variant displayed enhanced modulation of recovery, resembling zebrafish PRRT2 and significantly differing from human PRRT2 (Figure 3G). These findings indicate that the C-terminal region of zebrafish PRRT2 confers its greater ability to modulate recovery of Nav1.2 from the slow-inactivated state.

We next extended our analysis to other DspB family members, including *Trafficking regulator of GLUT4-1* (*trarg1*) and *Transmembrane protein 233* (*tmem233*), both of which encode proteins with C-terminal regions similar to that of PRRT2 (Figure 3-figure supplement 1A and B). Given the critical role of the PRRT2 C-terminus in modulating slow inactivation, we hypothesized that TRARG1 and TMEM233 might exert similar regulatory effects. Under heterologous expression conditions, both TRARG1 and TMEM233 facilitated the entry of Nav1.2 channels into the slow-inactivated state and delayed their recovery (Figure 3-figure supplement 1C and D). These effects were comparable to those observed with mouse PRRT2, although TMEM233 exhibited slightly weaker modulation than PRRT2 and TRARG1 (Figure 3-figure supplement 1C and D). Together, these results suggest that the role of PRRT2 in regulating Nav channel slow inactivation is evolutionarily conserved.

### PRRT2 promotes slow inactivation across multiple human Nav channel isoforms

In humans, the nine Nav channel isoforms (Nav1.1-1.9) share high sequence homology and similar structural features(Goldin, 2001). To determine whether PRRT2 modulates slow inactivation across different Nav channel isoforms, we selected Nav1.1, Nav1.4, Nav1.5, and Nav1.6, which are expressed in distinct tissues and cell types, and assessed the effects of human PRRT2 on their slow inactivation properties (Figures 4A-J).

**Figure 4.**
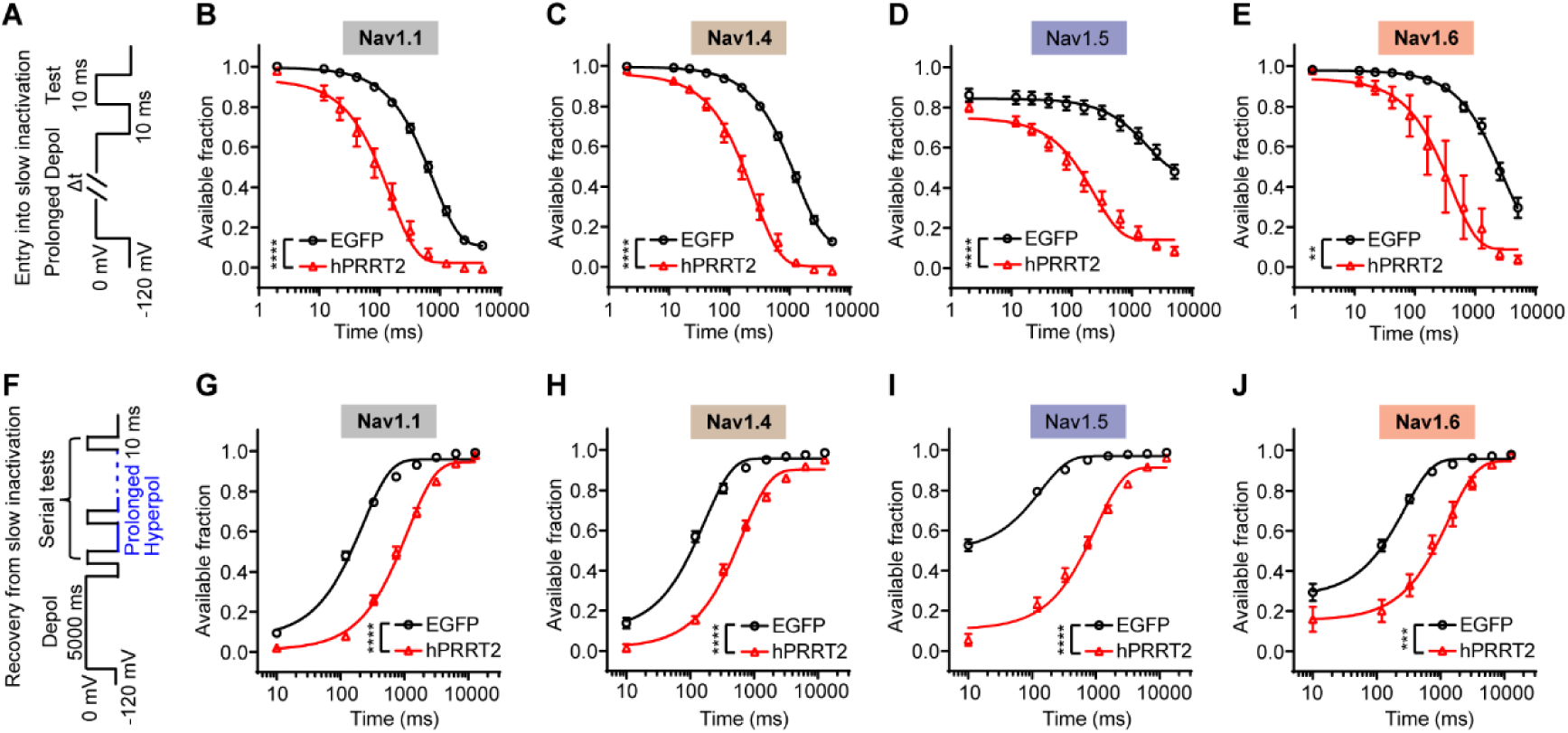
PRRT2 promotes slow inactivation across multiple human Nav channel isoforms. **(A)** Protocol for examining the entry of Nav channels into slow-inactivated state. **(B-E)** The effects of human PRRT2 (hPRRT2) on the entry of Nav1.1, Nav1.4, Nav1.5 and Nav1.6 channels into slow inactivation. In (B), analyses for Nav1.1 (EGFP: n = 10 cells, hPRRT2: n = 12 cells); In (C), analyses for Nav1.4 (EGFP: n = 11 cells, hPRRT2: n = 11 cells); In (D), analyses for Nav1.5 (EGFP: n = 17 cells, hPRRT2: n = 17 cells); In (E), analyses for Nav1.6 (EGFP: n = 5 cells, hPRRT2: n = 4 cells). **(F)** Protocol for examining the recovery of Nav channels from slow-inactivated state. **(G-J)** The effects of human PRRT2 on the recovery of Nav1.1, Nav1.4, Nav1.5 and Nav1.6 channels from slow-inactivated state induced by 5-s depolarization. In (G), analyses for Nav1.1 (EGFP: n = 10 cells, hPRRT2: n = 12 cells); In (H), analyses for Nav1.4 (EGFP: n = 11 cells, hPRRT2: n = 11 cells); In (I), analyses for Nav1.5 (EGFP: n = 16 cells, hPRRT2: n = 15 cells); In (J), analyses for Nav1.6 (EGFP: n = 5 cells, hPRRT2: n = 4 cells) Data are presented as mean ± s.e.m. The main effect of group was assessed using two-way ANOVAs (B-E and G-J). **P < 0.01, ***P < 0.001, ****P < 0.0001.

Nav1.1, Nav1.4, and Nav1.6 displayed similar kinetics in the development and recovery of slow inactivation (Figures 4B, C, E, G, H, and J). In contrast, Nav1.5 showed greater resistance to entering the slow-inactivated state (Figures 4D and I), consistent with previous findings(Richmond et al., 1998). Despite these isoform-specific differences, the heterologous expression of PRRT2 consistently promoted entry of Nav channels into the slow-inactivated state (Figures 4A-E) and delayed recovery from it (Figures 4F-J) across all four Nav isoforms tested. These results suggest that PRRT2 modulates slow inactivation through a common mechanism shared by multiple Nav channel isoforms, despite differences in their intrinsic slow-inactivation properties.

### Association between PRRT2 and Nav channels *in vitro*

PRRT2 may regulate slow inactivation through association with Nav channels. To examine the potential physical association between human PRRT2 and Nav1.2, we co-expressed PRRT2-HA and Flag-Nav1.2 in HEK293T cells and performed co-immunoprecipitation (Co-IP) assays (Figures 5A and B). As controls, cells were transfected with one of the two interacting proteins (either PRRT2-HA or Flag-Nav1.2) together with the empty vectors carrying the corresponding tag of the other protein (Figures 5A and B). Immunoprecipitation using an anti-Flag antibody to capture Flag-Nav1.2 successfully co-precipitated PRRT2 in the co-transfection group, but not in the control groups where either protein was expressed (Figure 5A). Conversely, immunoprecipitation with anti-HA antibody showed that Nav1.2 was specifically co-precipitated with PRRT2-HA in the co-transfection group but not in controls (Figure 5B). These results confirm that PRRT2 and Nav1.2 formed a protein complex *in vitro*.

**Figure 5.**
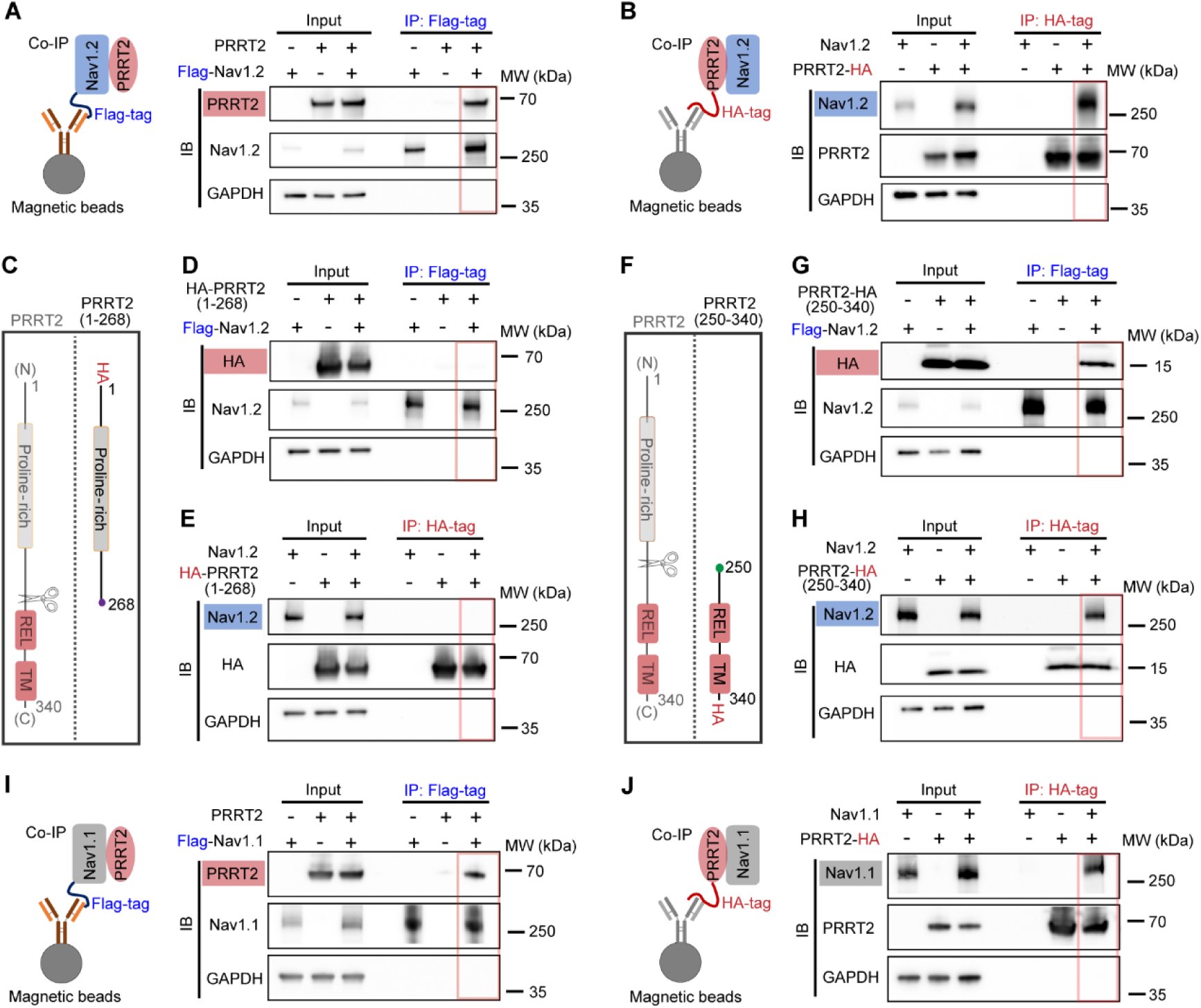
Association between PRRT2 and Nav channels in vitro. **(A and B)** Schematic diagram of the co-immunoprecipitation assay used to detect association between human PRRT2 and Nav1.2. Flag-tagged Nav1.2 (+), HA-tagged PRRT2 (+) and empty (-) vectors were transfected in HEK293T cells as indicated. The cell lysates were immunoprecipitated with anti-Flag (A) or anti-HA (B) magnetic beads, the captured proteins were analyzed by SDS-PAGE and immunoblotting. **(C)** Diagram for HA-tagged truncation of human PRRT2 (PRRT2(1–268)), in which the carboxyl terminus of PRRT2 was deleted. REL, re-entrant loop. TM, transmembrane domain. (D and E) Co-immunoprecipitation assay for potential association between Nav1.2 and PRRT2(1–268). The proteins immunoprecipitated by anti-Flag (D) or anti-HA (E) magnetic beads were analyzed by immunoblotting. Note that HA-tagged PRRT2(1–268) was detected by anti-HA antibody in (D and E). **(F)** Diagram for HA-tagged truncation of human PRRT2 (PRRT2(250–340)), in which the amino terminus of PRRT2 was deleted. **(G and H)** Co-immunoprecipitation assay for potential association between Nav1.2 and PRRT2(250–340). The proteins immunoprecipitated by anti-Flag (G) or anti-HA (H) were analyzed by immunoblotting. Note that HA-tagged PRRT2(250–340) was detected by anti-HA antibody in (G and H). **(I and J)** Co-immunoprecipitation assay for potential association between Nav1.1 and PRRT2. The proteins captured by anti-Flag (I) or anti-HA (J) magnetic beads were analyzed by immunoblotting. In (A, B, D, E, G, H, I and J), blots shown are representative of at least three independent co-immunoprecipitation experiments with similar results. Red rectangles indicate the immunoblotting of bait and potential prey proteins in co-immunoprecipitation. IB, immunoblot. IP, immunoprecipitation. MW, molecular weight.

To identify the Nav1.2-associating region within human PRRT2, we generated truncated constructs encoding either the N-terminus (residues 1–268) or C-terminus (residues 250–340) of PRRT2. Using the same Co-IP approach, we found that PRRT2(1–268), which contains the intracellular proline-rich domain, failed to associate with Nav1.2 (Figures 5C-E). In contrast, the PRRT2(250–340), which includes the re-entrant loop and transmembrane domain, was able to interact with Nav1.2 (Figures 5F-H). These results indicate that the C-terminal region of PRRT2 is necessary and sufficient, in this assay, for association with Nav1.2 channels. Given that PRRT2 modulates slow inactivation across multiple Nav isoforms, we hypothesized that it might also associate with other Nav isoforms, such as Nav1.1. To test this, we co-expressed PRRT2-HA and Flag-Nav1.1 in HEK293T cells and performed Co-IP assays. Under these conditions, PRRT2 was found to associate with Nav1.1 (Figures 5I and J), similar to the association observed with Nav1.2 (Figures 5A and B).

These findings contrast with previous reports, which suggested that PRRT2 preferentially interacted with Nav1.2 over Nav1.1 *in vitro*(Franchi et al., 2023; Fruscione et al., 2018; Valente et al., 2023). This discrepancy may stem from differences in the experimental conditions used to assess protein-protein interactions. For example, in our study, Nav1.1 and PRRT2 were co-expressed in HEK293T cells, and cells were solubilized using the detergent n-Dodecyl β-D-maltoside (DDM). Variations in experimental conditions could influence the stability of the PRRT2-Nav channel complex and the efficiency of the Co-IP assay.

Collectively, our findings suggest that PRRT2 associates with Nav channels through its membrane-associated C-terminal domains, forming molecular complexes *in vitro*.

### PRRT2 forms a molecular complex with Nav1.2 in the mouse brain

To extend our findings from the heterologous expression system to an *in vivo* context, we first investigated whether PRRT2 forms a protein complex with Nav channels in brain tissue. Due to the limited efficacy of available antibodies for immunoprecipitating native PRRT2 or Nav1.2 from brain lysates, we employed a tag-based capture strategy, which relies on the appropriate insertion of a commonly used epitope tag into the target protein.

We generated *Prrt2-V5* knock-in mice via CRISPR/Cas9 technology(Yang, Wang, & Jaenisch, 2014), in which a V5 epitope tag was inserted at the C-terminus of PRRT2 (Figures 6A and B). Notably, compared with wild-type mice, PRRT2 protein levels in tissue lysates were markedly reduced in *Prrt2-V5* knock-in mice (Figure 6C-D). These mice displayed susceptibility to dystonia attacks, resembling the phenotype previously observed in *Prrt2*-mutant mice(Wu et al., 2025), but with lower penetrance (Figure 6-figure supplement 1A-C). Despite the reduced PRRT2 expression, Nav1.2 expression remained unchanged (Figure 6E), and co-immunoprecipitation assays remained effective in the *Prrt2-V5* knock-in mice. Using anti-V5 nanobody-conjugated magnetic beads, PRRT2 was effectively immunoprecipitated from membrane lysates of brain tissue (Figure 6C). A notable amount of Nav1.2 was co-immunoprecipitated in the *Prrt2-V5* knock-in group, whereas only trace levels were detected in wild-type controls (Figures 6C and F). In contrast, ATP1B2, another membrane protein, was not co-immunoprecipitated in the *Prrt2-V5* knock-in group (Figures 6C and F), supporting the specificity of the association between PRRT2 and Nav1.2. Together, although the *V5* knock-in allele is hypomorphic, these results provide *in vivo* evidence that PRRT2 forms a molecular complex with Nav1.2 channels in the mouse brain.

**Figure 6.**
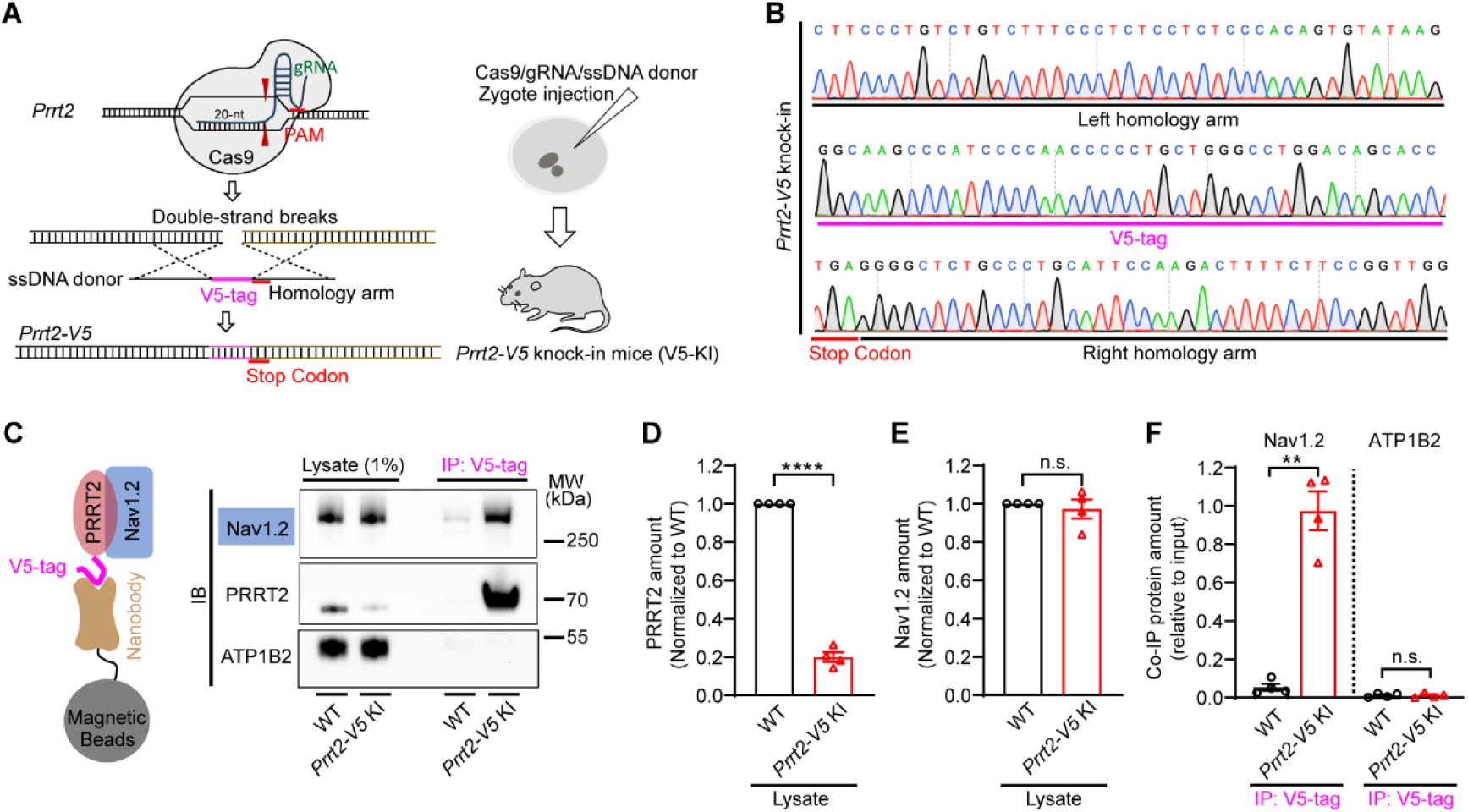
PRRT2 forms a molecular complex with Nav1.2 in the mouse brain. **(A)** Schematic showing the generation of Prrt2-V5 knock-in (V5-KI) mice using Cas9 technology. V5-tag sequence was inserted precisely at the end of Prrt2 gene. PAM, protospacer-adjacent motif (NGG). ssDNA, single-strand DNA. **(B)** Sanger sequencing around sgRNA targeting site in Prrt2-V5 knock-in mouse. The V5-tag, stop codon and homologous arms were denoted. **(C)** Co-immunoprecipitation assay for potential interaction between Nav1.2 and PRRT2-V5 in the brain tissue. The proteins immunoprecipitated by anti-V5 nanobody were analyzed by immunoblotting. Blots shown are representative of four independent co-immunoprecipitation experiments. IB, immunoblot. IP, immunoprecipitation. MW, molecular weight. WT, wild-type. KI, knock-in. **(D-F)** Quantification of the density of protein bands for PRRT2 (D) and Nav1.2 (E) in lysate, and for Nav1.2 and ATP1B2 in co-immunoprecipitation experiments (F) (n = 4 mice for each group). Data are presented as mean ± s.e.m. In (D-F), two-tailed, paired Student’s t-test was used for determining statistical significance. **P < 0.01, ****P < 0.0001, n.s., not significant.

### PRRT2 deficiency impairs the regulation of Nav channel slow inactivation in cortical neuronal axons

Given the dense distribution of both PRRT2 and Nav channels in the cerebral and cerebellar cortices(Yamano, Miyazaki, & Nukina, 2022), we next investigated whether PRRT2 regulates Nav channel slow inactivation in cortical neurons. Cortical slices were prepared from wild-type and *Prrt2*-mutant mice, the latter lacking PRRT2 expression(Tan et al., 2018). Unlike the relatively simple morphology of HEK293T cells, pyramidal neurons have complex arborizations, which present space-clamp limitations during whole-cell voltage-clamp recordings(Bar-Yehuda & Korngreen, 2008). To mitigate these issues, we followed established protocols to isolate axonal blebs from cortical pyramidal neurons(Hu & Shu, 2012). These spherical, membrane-bound blebs form at the cutting surface of cortical slices and are readily visualized (Figure 7A). Sodium currents recorded from axon blebs were voltage-dependent and sensitive to tetrodotoxin (TTX) (Figure 7A), confirming that they were primarily mediated by Nav channels(Hu & Shu, 2012).

**Figure 7.**
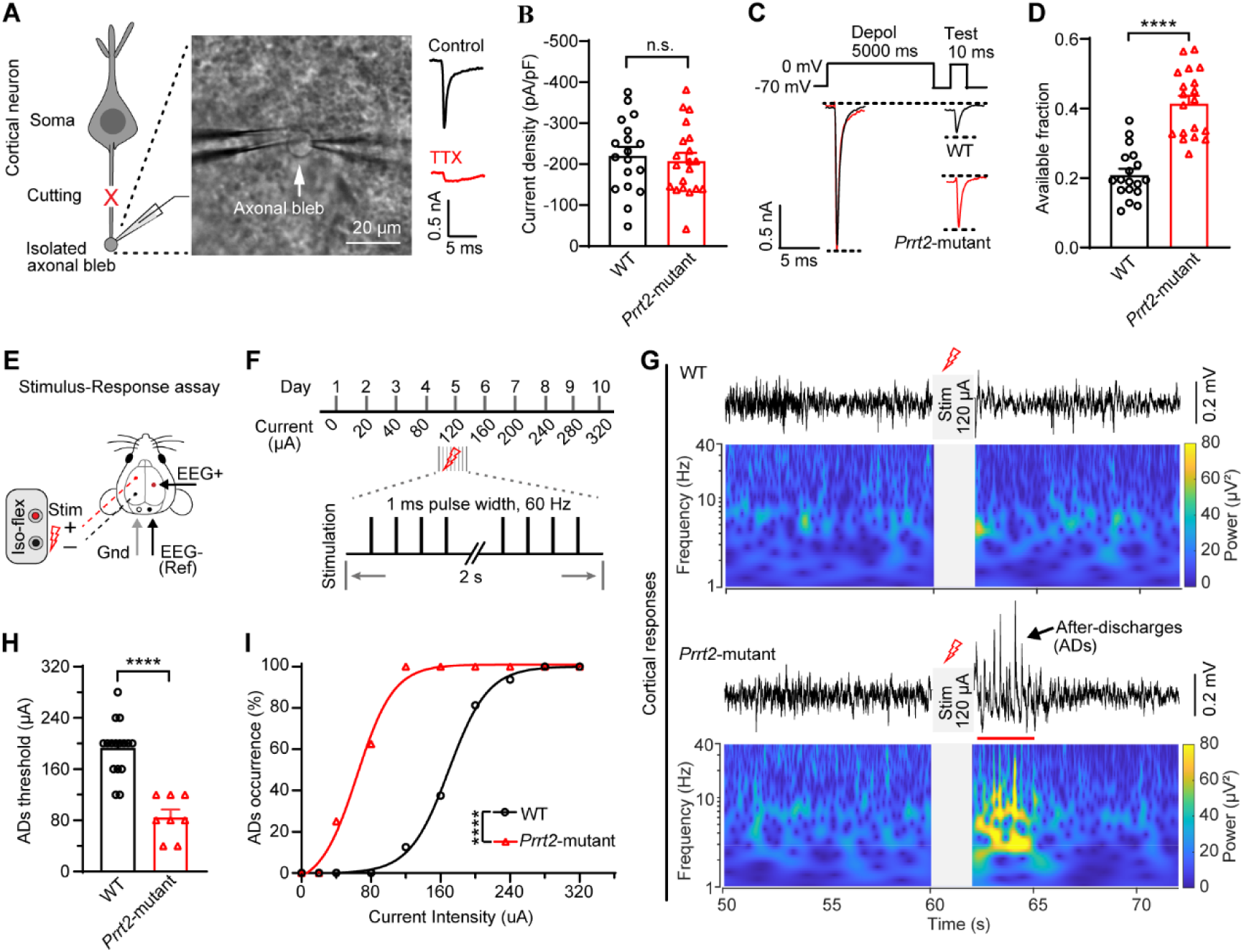
PRRT2 deficiency impairs the regulation of Nav channel slow inactivation and neuronal resilience in mice. **(A)** Schematic showing the isolated axonal bleb recording in cortical neuron (left) and the representative brightfield image for the axonal bleb in brain slice (middle). The arrow indicates an axonal bleb. Scale bar, 20 μm. The representative sodium current traces recorded in axonal blebs (right). Scale bar, 0.5 nA, 5 ms. TTX, tetrodotoxin. **(B)** Sodium current density recorded in isolated axonal blebs from wild-type (WT) and Prrt2-mutant mice (WT: n = 18 blebs from 7 mice, Prrt2-mutant: n = 20 blebs from 8 mice). Sodium currents were measured with a brief depolarization to 0 mV preceded by a 200-ms hyperpolarizing prepulse to -110 mV. **(C)** Protocol for assessing Nav channel slow-inactivation (upper) and the representative traces of the sodium currents evoked by conditioning depolarization pulse and test pulses (bottom). Scale bar, 0.5 nA, 5 ms. **(D)** Fraction of available Nav channels after 5-s depolarization (WT: n = 18 blebs from 7 mice, Prrt2-mutant: n = 19 blebs from 8 mice). **(E)** Schematic illustration of the cortical electrostimulation and EEG recording in mice. Ref, reference electrode. Gnd, ground electrode. Stim, Stimulation. **(F)** Illustration showing the experimental schedule and the stimulation parameters. Stimulation was delivered once daily with stepwise increases in current intensity. **(G)** Representative traces and power of EEG signals in wild-type and Prrt2-mutant mice before and after 2-s cortical stimulation. After-discharges were indicated by the arrow and red line. Scale bar: 0.2 mV. Electrical stimulation (120 µA) was applied during 60–62 s period, where the stimulus-induced artifacts were removed prior to analysis. **(H)** Threshold of electrical stimulation to induce after-discharges in WT and Prrt2-mutant mice (WT: n = 16 mice, Prrt2-mutant: n = 8 mice). **(I)** Percentage of after-discharge occurrence in WT and Prrt2-mutant mice after electrical stimulation (WT: n = 16, Prrt2-mutant: n = 8). Data are presented as mean ± s.e.m. Two-tailed, unpaired Student’s t-test was used in (B and D) and two-tailed, unpaired Mann-Whitney test was used in (H). In (I), the main effect of group was assessed using two-way ANO-VAs. ****P < 0.0001, n.s., not significant.

PRRT2 deficiency did not affect Nav1.2 protein expression in forebrain tissues (Figure 7-figure supplement 1A and B), nor did it alter the sodium current density in cortical axonal blebs when measured with a brief depolarization to 0 mV preceded by a 200-ms hyperpolarizing prepulse to -110 mV, a protocol designed to relieve inactivation before current elicitation (Figure 7B). Because prolonged voltage-clamp recordings from isolated axonal blebs are technically challenging, we adopted a simplified protocol to assess Nav channel slow inactivation, using a holding potential of -70 mV to approximate the resting membrane potential of cortical neurons. In this protocol, a single 5-s depolarization to 0 mV was applied, and the fraction of available Nav channels was determined after a 10-ms hyperpolarization (Figure 7C).

Approximately 20% of available sodium current remained in wild-type blebs, compared to ∼40% in *Prrt2*-mutant blebs (Figures 7C and D), indicating that during sustained depolarization, fewer Nav channels in the *Prrt2*-mutant group entered the slow-inactivated state. Notably, because the proportion of channels entering slow inactivation differed substantially between wild-type and *Prrt2*-mutant groups, direct comparison of recovery kinetics was not feasible and was therefore not performed. Nevertheless, these findings provide direct evidence that endogenous PRRT2 is required for effective regulation of Nav channel slow inactivation in cortical neurons.

### PRRT2 deficiency compromises cortical resilience in awake mice

Excitatory neurons exhibit intrinsic adaptation, whereby their responses progressively decline during sustained or repetitive activation(Fleidervish, Friedman, & Gutnick, 1996; Kim, Owen, Holmes, & Grover, 2012). We therefore hypothesized that PRRT2 may contribute to this process by promoting Nav channel slow inactivation.

To test this idea, we applied a repetitive stimulation protocol in the corpus callosum, a white-matter region of the brain enriched in both PRRT2 and Nav channels. Compound action potentials, recorded with a glass electrode positioned 150 μm from the stimulation site, were used as a functional readout related to Nav channel availability during repetitive activity (Figure 7-figure supplement 2A). Compared with wild-type mice, *Prrt2*-mutant mice exhibited a smaller decline in compound action potential amplitude during high-frequency stimulation (20 Hz), but not during low-frequency stimulation (1 Hz), indicating reduced adaptation (Figure 7-figure supplement 2B and C). This pattern is consistent with impaired slow inactivation during repetitive activity.

Neuronal adaptation is important for limiting excitability during episodes of aberrant network hyperactivity (Fleidervish et al., 1996; Zang, Marder, & Marom, 2023). To further determine whether PRRT2-dependent impairment of neuronal adaptation contributes to hyperexcitability *in vivo*, we next assessed cortical resilience in awake mice, a measure of the resistance of the cerebral cortex to pathological perturbations. We used a stimulation-response paradigm in which stimulating electrodes were implanted into the left cerebral cortex, targeting both the sensory and visual cortices, while EEG recording electrodes were placed in the contralateral hemisphere, which receives projections from the stimulated region (Figure 7E). Pulsed electrical stimulation (1-ms pulse width at 60 Hz for 2 s) was delivered to the cortex of mice once daily with stepwise increases in current intensity, and the minimal current required to evoke after-discharges (ADs) in the contralateral cortex was determined in each animal (Figure 7F). The threshold current required to induce after-discharges was used as a functional indicator of cortical resilience to aberrant excitatory inputs(Blume, Jones, & Pathak, 2004).

*Prrt2*-mutant mice exhibited significantly lower stimulation thresholds for evoking after-discharges compared to wild-type controls (Figures 7G and H). Specifically, stimulation at an average intensity of 85 µA reliably induced after-discharges in *Prrt2*-mutant mice, whereas a significantly higher intensity (190 μA) was required to elicit similar responses in wild-type mice (Figures 7H and I), indicating that PRRT2 contributes importantly to maintaining cortical resilience in response to excitatory perturbations.

## Discussion

In this study, we identified PRRT2 as a physiological auxiliary factor of Nav channel slow inactivation. We demonstrate that PRRT2 exerts a dual action by both facilitating the transition of Nav channels into a slow-inactivated state and delaying their recovery. Importantly, this regulatory action is evolutionarily conserved, and requires the integrity of the PRRT2 C-terminal region. Using both *Prrt2-V5* knock-in and *Prrt2*-mutant mouse models, we provide compelling *in vivo* evidence that PRRT2 forms a molecular complex with Nav channels and modulates their slow inactivation, thereby contributing to neuronal adaptation and cortical resilience to perturbations.

In eukaryotic Nav channels, ion permeability is regulated through at least four distinct and coordinated gating mechanisms: voltage-gated activation and deactivation, fast inactivation, and slow inactivation(Catterall, 2023). Slow inactivation, an evolutionarily conserved intrinsic property of Nav channels, is mediated by conformational rearrangements and has distinct molecular determinants from those governing fast inactivation(Payandeh, 2018; Silva, 2014; Vilin & Ruben, 2001).

The role of PRRT2 in the regulation of Nav channel slow inactivation appears to be unique compared to known regulators such as SCN1B and FHFs. The modulatory effects of the SCN1B on Nav channel inactivation were initially observed in *Xenopus* oocytes(Wallner, Weigl, Meera, & Lotan, 1993), where it was found to influence sodium currents. Subsequent studies proposed that the SCN1B impedes slow inactivation, a hypothesis later confirmed in mammalian cells co-expressing Nav1.4 and SCN1B(Webb et al., 2009). However, its regulatory effects on slow inactivation varied across Nav isoforms(He & Soderlund, 2014; Makita, Bennett, & George, 1996; Vilin et al., 1999; Xu et al., 2007). In this study, we observed little effect of SCN1B on the onset and recovery of slow inactivation in Nav1.2, consistent with previous findings(Xu et al., 2007). In contrast to SCN1B, PRRT2 exerts a more consistent effect across Nav isoforms, enhancing slow inactivation regardless of the specific Nav subtype. FHF proteins (FHF1-4, also known as FGF11-14) constitute a subfamily of FGFs that lack a secretion signal and function distinctly from traditional FGFs(Olsen et al., 2003). Previous studies demonstrated that FHFs mediate a rapid-onset, long-term inactivation, which differs mechanically from traditional slow inactivation(Dover et al., 2010). Unlike the FHF2A-mediated long-term inactivation, which can be triggered by a brief 2-ms depolarization, PRRT2 requires a longer depolarization to produce a detectable effect on the onset of slow inactivation of Nav channels. Additionally, PRRT2 significantly delays recovery of Nav channels from the slow-inactivated state induced by 5-second depolarization, whereas FHF2A has little effect on this process. These findings suggest that PRRT2 and FHF2A modulate Nav channel inactivation through different mechanisms.

Previous studies in heterologous overexpression systems have shown that PRRT2 can influence Nav channel trafficking and surface expression(Fruscione et al., 2018; Valente et al., 2023), raising the possibility that the observed effects on slow inactivation are secondary to altered channel abundance or localization. However, slow inactivation develops on a timescale of hundreds of milliseconds to seconds, whereas detectable changes in Nav channel trafficking and surface abundance generally occur over much longer intervals (minutes to hours) (Freal et al., 2023; Higerd-Rusli et al., 2023). These distinct temporal profiles argue against trafficking as the primary basis for the effects of PRRT2 on Nav channel slow inactivation described here, although direct quantification of dynamic changes in Nav channel surface expression will be required to fully exclude such a contribution(Liu, Wang, Pitt, & Liu, 2022; Tyagi, Higerd-Rusli, Akin, Waxman, & Dib-Hajj, 2025).

Our truncation analysis showed that the C-terminal region of PRRT2, which contains the re-entrant loop and transmembrane domain, is sufficient to reproduce the enhancement of slow inactivation. This finding suggests that PRRT2 modulates slow inactivation through a membrane-embedded or membrane-proximal mechanism, rather than through its intracellular pro-line-rich N-terminus. Current models of Nav channel slow inactivation propose that entry into the slow-inactivated state involves coupled conformational changes in both the voltage-sensing domains and the pore region(Catterall, Gamal El-Din, & Wisedchaisri, 2024; Silva, 2014), including contributions from the external pore mouth(Balser et al., 1996; Xiong et al., 2006), the selectivity filter(Chatterjee et al., 2018; Todt, Dudley, Kyle, French, & Fozzard, 1999), the S6 segments(Chancey, Shockett, & O’Reilly, 2007; Y. Chen, Yu, Surmeier, Scheuer, & Catterall, 2006; S. Y. Wang & Wang, 1997), and regions encompassing S4, S4-S5 linker, and S5 (Bendahhou, Cummins, Kula, Fu, & Ptacek, 2002; Cummins & Sigworth, 1996; Hayward et al., 1999; Mitrovic, George, & Horn, 2000; D. W. Wang, Makita, Kitabatake, Balser, & George, 2000). Within this framework, the membrane-associated C-terminal region of PRRT2 may enhance slow inactivation by stabilizing one or both of these conformational rearrangements, although the precise structural basis of this effect remains to be determined.

In mammals, Nav channel isoforms (Nav1.1–1.9) are distributed across the nervous system, skeletal muscle, heart and secretory glands(Cusdin, Clare, & Jackson, 2008). Although we have shown that PRRT2 exerts a consistent effect on slow inactivation across various Nav channel isoforms *in vitro*, the regulation of Nav channel slow inactivation is likely to be more complex under physiological conditions. PRRT2 is predominantly expressed in the nervous system(W. J. Chen et al., 2011), with enrichment in excitatory neurons(Tan et al., 2018), making it unlikely to directly modulate Nav1.4 and Nav1.5 channels, which are primarily expressed in skeletal muscle and cardiac tissue, respectively(Cusdin et al., 2008). A similar rationale applies to Nav1.1, which is largely confined to inhibitory interneurons(Yu et al., 2006), where PRRT2 expression is minimal or absent. These observations suggest that the functional consequence of PRRT2 depend on the Nav isoform composition and cellular context of each tissue.

The partial mismatch between PRRT2 and Nav channel expression *in vivo* raises an important question about how Nav channel slow inactivation is regulated in cells that do not express PRRT2. One possibility is the expression of alternative regulators, such as TRARG1 or TMEM233, both of which belong to the DspB family and are expressed in tissues distinct from PRRT2(Ehrlich, Lacey, & Ehrlich, 2020; Oort, Warden, Baumann, Knotts, & Adams, 2007; Santana-Varela et al., 2021; Shibata et al., 2007). These proteins may serve analogous functions in modulating Nav channel slow inactivation. Another possibility is that certain PRRT2-negative cells have a reduced requirement for Nav channel slow inactivation modulation due to their intrinsic rhythmic firing properties. For instance, in the cerebellar cortex, the PRRT2 is absent in Purkinje cells, which exhibit repetitive firing(Lu et al., 2021). In such rhythmic firing cells, strong enhancement of Nav channel slow inactivation may be undesirable, as excessive accumulation of inactivated Nav channels could disrupt their regular firing patterns. Thus, the broad isoform activity of the PRRT2 should be considered in any future attempt to manipulate PRRT2 function therapeutically. Notably, Nav channel slow inactivation represents only one of several mechanisms that regulate neuronal excitability. In both PRRT2-positive and PRRT2-negative neurons, excitability may also be fine-tuned through alternative mechanisms, including regulation of potassium channels such as Kv7.2/Kv7.3 and additional ion channel- or signaling-dependent pathways(Debanne, Inglebert, & Russier, 2019; Jones, Gamper, & Gao, 2021). These distinct mechanisms likely operate in a coordinated manner to maintain neuronal excitability, with PRRT2-dependent regulation of Nav channel slow inactivation contributing serving as one component of this broader homeostatic framework(Marom & Marder, 2023).

Given the critical role of PRRT2 in regulating the slow inactivation of Nav1.2 and Nav1.6 channels, both of which are highly expressed in cortical neurons, PRRT2 deficiency may alter neuronal excitability and neuronal responses to prolonged or repetitive activity, thereby contributing to central nervous system dysfunction. Evidence from cultured neurons, animal models, and human genetics is broadly consistent with this view. Studies using human induced pluripotent stem cells (iPSC)-derived excitatory neurons have shown that homozygous *PRRT2* mutation increases sodium currents and neuronal excitability(Fruscione et al., 2018). In mouse models, loss-of-function mutation or knockout of PRRT2 in cerebellar granule cells increases intrinsic excitability and facilitates spreading depolarization in the cerebellar cortex(Binda, Valente, Marte, Baldelli, & Benfenati, 2021; Lu et al., 2021), whereas in the forebrain, PRRT2 deficiency impairs Nav channel slow inactivation in cortical neurons and reduces resistance to pathological perturbation in awake mice. In addition, *PRRT2* mutations in humans are associated with infantile convulsions(Lee et al., 2012), paroxysmal dyskinesia(W. J. Chen et al., 2011; J. L. Wang et al., 2011), and hemiplegic migraine(Riant et al., 2022), all disorders linked to abnormal neuronal network excitability.

Although the present study supports a functional link between PRRT2-dependent regulation of Nav channel slow inactivation and cortical resilience, PRRT2 may influence cortical excitability and stability through additional Nav-dependent mechanisms, including reduced current density and shifts in the voltage dependence of channel inactivation(Fruscione et al., 2018; Lu et al., 2021; Valente et al., 2023). Notably, because PRRT2 facilitates Nav channel entry into slow-inactivated states both from closed states and from open states during prolonged depolarization, some of these previously reported effects may partly reflect enhanced slow inactivation and the resulting reduction in Nav channel availability. In addition to these Nav-dependent mechanisms, previous studies have shown that PRRT2 also regulates synaptic vesicle cycling(Coleman et al., 2018; Tan et al., 2018; Valente et al., 2016) and presynaptic surface expression of Cav2.1 channels(Ferrante et al., 2021). These effects are also expected to influence neurotransmitter release and, consequently, neuronal and network excitability, and may therefore contribute to cortical resilience.

Together, the identification of PRRT2 as an auxiliary factor regulating Nav channel slow inactivation provides new insight into the fine-tuning of Nav channel function and offers a mechanistic framework for understanding how PRRT2 deficiency may contribute to episodic neurological disorders associated with excitability dysregulation.

### Limitations of the study

While our study identifies the functional domain of PRRT2 and provides biological evidence that PRRT2 associates with Nav1.2, the precise binding site on the Nav channel that mediates PRRT2-dependent regulation of slow inactivation remains undefined. Direct structural and biochemical mapping of the PRRT2-Nav interface, including targeted mutagenesis, crosslinking, and structural determination, will be necessary to elucidate the molecular basis of this interaction and its effect on channel gating. Cryo-electron microscopy (cryo-EM) represents a promising approach for resolving the structural details of the Nav-PRRT2 complex, particularly in light of recent advancements in determining high-resolution structures of various Nav channel isoforms and their complexes with auxiliary factors(Jiang et al., 2020; Pan et al., 2019; Pan et al., 2021).

## Materials and Methods

### Molecular cloning

Mammalian expression constructs were generated using the pCAGIG backbone (Addgene, 11159), which contains an internal ribosome entry site (IRES) followed by EGFP(Matsuda & Cepko, 2004). Coding sequences for full-length or truncated proteins of interest (POIs) were inserted in the pCAGIG vector using a seamless cloning method (Beyotime, D7010). The POIs included human PRRT2 (hPRRT2), hPRRT2(1–268), hPRRT2(250–340), mouse PRRT2 (mPRRT2), mPRRT2(222–346), mPRRT2(256–346), mPRRT2(1–266), zebrafish PRRT2 (zfPRRT2), mouse SCN1B, mouse FHF2A, mouse TRARG1 and mouse TMEM233. A PRRT2 chimeric construct was generated by fusing the N-terminal fragment of human PRRT2 (hPRRT2(1–222)) with the C-terminal region of zebrafish PRRT2 (zPRRT2(110–226)), and cloning the resulting sequence into pCAGIG vector. Plasmids encoding human PRRT2(W. J. Chen et al., 2011) were kindly provided by Dr. Zhi-Ying Wu’s Laboratory (Zhejiang University). Plasmids of pCAG-Flag-Nav1.2(Pan et al., 2019) and pCAG-Flag-Nav1.1(Pan et al., 2021) were provided by Huai-Zong Shen’s Laboratory (Westlake University). All constructs generated in this study were verified by Sanger sequencing (BioSune, Boshang Biology Technology, Shanghai).

### Cell culture and heterologous expression

HEK293T cell lines stably expressing human Nav1.2 or Nav1.6 and CHO cell lines stably expressing human Nav1.1, Nav1.4 or Nav1.5 were obtained from the laboratory of ICE Bioscience Inc. Wild-type HEK293T cells (SCSP-502) used for transient transfection were obtained from Cell Bank/Stem Cell Bank of Chinese Academy of Sciences. All cells were cultured in Dulbecco’s modified Eagle medium (DMEM; Gibco, 11995065) supplemented with 10% fetal bovine serum (FBS; Gibco,10099141) and penicillin (50 U/mL)-streptomycin (50 μg/mL) (Gibco,15140163), and maintained at 37 °C in a humidified incubator with 5% CO_2_. The HEK293T cell line used in this study is of female origin and tested negative for mycoplasma contamination.

For heterologous protein expression in whole-cell voltage-clamp recording, 2 μg of plasmid DNA was transfected into HEK293T or CHO cells using Lipofectamine 2000 (ThermoFisher, 11668-019) according to the manufacturer’s instructions. For co-immunoprecipitation assays, 2–3 μg of total plasmid DNA (encoding two proteins of interest) was used for co-transfection. Four hours after the transfection, the medium was replaced with fresh growth medium, and cells were maintained for an additional 20 hours before subsequent experiments.

### Animals

*Prrt2*-mutant mice, carrying a premature stop codon in exon 2 of the *Prrt2* gene(Tan et al., 2018), were used in electrophysiological recording experiments.

*Prrt2*-*V5* knock-in mice, used for co-immunoprecipitation (Co-IP) experiments, were generated in this study via CRISPR/Cas9 technology(Yang et al., 2014). Briefly, single-guide RNA (sgRNA: 5’-TCTCCCACAGTGTATAAGTG-3’) targeting the location nearby the stop codon of *Prrt2* gene allele, together with a single-stranded DNA donor template encoding a V5 tag flanked by 53-bp homology arms (ssDNA: 5’-CTTCCCTGTC TGTCTTTCCC TCTCCTCTCC CACAGTGTAT AAGGGCAAGC CCATCCCCAA CCCCCTGCTG GGCCTGGACA GCAC-CTGAGG GGCTCTGCCC TGCATTCCAA GACTTTTCTT CCTGTTGG-3’) and Cas9 mRNA, were microinjected into zygotes from fertilized C57BL/6J mice. The injected zygotes were implanted into pseudopregnant female mice. Offspring were screened for correct insertion of the V5 tag immediately upstream of the *Prrt2* stop codon by PCR, and the insertion was subsequently verified by Sanger sequencing. Positive founder (F0) mice were crossed with C57BL/6J mice to establish the *Prrt2*-*V5* knock-in line.

Both *Prrt2*-mutant and *Prrt2*-*V5* lines were backcrossed onto the C57BL/6J background for more than 10 generations. C57BL/6J mice used for breeding were obtained from the Shanghai Laboratory Animal Center (SLAC), Chinese Academy of Science.

Mice were housed in a 12-h light-dark cycle (light on at 7:00 a.m.) with ad libitum access to food and water. All procedures involving animals were approved by the Animal Care and Use Committee of the Center for Excellence in Brain Science and Intelligence Technology, Chinese Academy of Sciences (Approval number, NA-009-2022). Both male and female mice were used in experiments.

### Whole-cell voltage clamp recording for sodium currents in cell lines

Sodium currents were recorded from HEK293T or CHO cell lines stably expressing individual human Nav channel isoforms (ICE Bioscience Inc.). Cells were transfected with plasmids as described in the main text. Twenty-four hours after transfection, cells were trypsinized and re-plated at lower density onto poly-D-lysine-coated coverslips. EGFP-positive cells were selected for recording 24 hours later.

Whole-cell voltage-clamp recordings were conducted at room temperature (23-24 °C). The bath solution contained (in mM): 50 NaCl, 90 choline chloride, 3.5 KCl, 1 MgCl_2_, 2 CaCl_2_, 10 D-Glucose, 1.25 NaH_2_PO_4_, and 10 HEPES (pH adjusted to 7.4 with NaOH). The internal pipette solution contained (in mM): 50 CsCl, 60 CsF, 10 NaCl, 20 EGTA, 10 HEPES (pH adjusted to 7.2 with CsOH). The osmolarity of the external and internal solutions was adjusted to 300 and 290 mOsmol/kg, respectively.

Pipette electrodes were pulled from borosilicate glass capillary (Sutter Instruments, BF150-86-10) using a horizontal puller (P-97, Sutter Instruments). The pipette resistance was approximately 2 MΩ when filled with internal pipette solution. After whole-cell configuration was achieved, cells were allowed to equilibrate for 5 minutes prior to recording. Series resistance was compensated by 80% to minimize voltage errors. Currents were recorded with an amplifier (HEKA, EPC-10), sampled at 10 kHz, and acquired using Patchmaster (HEKA, v2x92).

### Fast inactivation assay for Nav channels

To assess the fast-inactivation kinetics of Nav channels, sodium currents were evoked by 20-ms depolarizing steps to test potentials between -20 and +30 mV in 10-mV increments, from a holding potential of -120 mV. The decay time constant (tau) was obtained by mono-exponential fitting of the current trace.

Recovery from fast inactivation was measured using a two-pulse protocol, in which a 10-ms depolarizing pulse to 0 mV (P1) was followed by a variable-duration recovery period at -120 mV and a second 10-ms test pulse at 0 mV (P2). Recovery was quantified as the normalized peak current of P2 relative to P1.

### Slow inactivation assay for Nav channels

To evaluate the development of slow inactivation of Nav channels, a series of conditioning prepulses of increasing duration at 0 mV was applied, followed by a brief 10-ms recovery at -120 mV and a 10-ms test pulse at 0 mV. Inter-sweep intervals were set at 15 s. Available fractions of Nav channels were assessed by normalizing the peak currents in the test pulse to that of the conditioning pulse.

For slow inactivation recovery, a 5-s conditioning pulse at 0 mV was followed by variable recovery durations at -120 mV, then a 10-ms test pulse at 0 mV. The inter-sweep interval was 15 s.

In experiments assessing the effect of zebrafish PRRT2 on slow inactivation recovery, a modified protocol was used to avoid cumulative inactivation. Within a single sweep, a 5-s depolarization was followed by a series of 10-ms test pulses to 0 mV, each separated by progressively longer recovery periods at -120 mV(Webb et al., 2009).

In this study, we chose 0 mV as the primary conditioning potential because it is widely used in conventional slow inactivation protocols and induces slow inactivation more robustly than more negative conditioning voltages, such as -70 mV.

### Steady-state slow inactivation assay for Nav channels

To measure steady-state slow inactivation, 10-s conditioning pulses ranging from -110 to -10 mV (in 10 mV increments) were applied, followed by a 10-ms hyperpolarizing step to -120 mV and subsequently a 5-ms test pulse to 0 mV. The peak test current was normalized to the maximum sodium current recorded across all conditions in the protocol and reported as I/I_max_.

### Slice preparation

Mice (postnatal day 14-20) were anesthetized with 4% isoflurane. The brain tissues were rapidly extracted and immersed in ice-cold sucrose-based cutting solution containing (in mM): 2.5 KCl, 1.25 NaH_2_PO_4_, 26 NaHCO_3_, 10 MgSO_4_, 0.5 CaCl_2_, 10 D-Glucose, 205 sucrose, 1 sodium pyruvate, 1.3 sodium L-ascorbate. Coronal cortical slices (250 μm) were prepared using a vibratome (Leica, VT1200S).

Slices were then transferred immediately to an incubation beaker containing aerated artificial cerebrospinal fluid (ACSF) composed of (in mM): 126 NaCl, 2.5 KCl, 2 MgSO_4_, 2 CaCl_2_, 26 NaHCO_3_, 1.25 NaH_2_PO_4_ and 25 dextrose (pH 7.4, 315 mOsm). Slices were incubated at 35 °C for 30 min, then maintained in the same solution at room temperature until use. The oxygenated ACSF was equilibrated with 95% O_2_ and 5% CO_2_.

### Electrical stimulation and field recording in the corpus callosum

Coronal slices were transferred to a recording chamber. A stimulation electrode (125 µm in diameter, concentric bipolar electrode, FHC, USA) was placed on the corpus callosum. The Ag-AgCl recording electrode (2-3 MΩ) was located at a site 150 µm away from the stimulation electrode along the corpus callosum. Stimulating pulses (0.1 ms pulse width, 100 µA) were delivered at 1 Hz for 100 seconds or at 20 Hz for 5 seconds. Compound action potentials were recorded as a functional readout related to Nav channel availability.

### Electrophysiological recording in isolated axonal blebs

Slices were transferred to a recording chamber and stabilized with a platinum anchor grid during recording. Resealed axonal ends (“blebs”) in the cerebral cortex were visualized using an upright infrared differential interference contrast (IR-DIC) microscope. Individual blebs were mechanically isolated by sweeping a cutting pipette beneath it, and subsequently patched with glass electrodes (resistance ∼7 MΩ).

For sodium current recordings, pipettes were filled with a potassium gluconate-based internal solution containing (in mM): 130 K-gluconate, 1 MgCl_2_, 11 EGTA, 1 CaCl_2_, 1 KCl, 2 Mg-ATP, 0.3 Na-GTP, 10 HEPES (pH adjusted to 7.2, ∼290 mOsm), supplemented with 2 mM TEA-Cl. The external ACSF solution was supplemented with 5 mM 4-Aminopydine (4-AP) and 200 μM CdCl_2_ to block K^+^ and Ca^2+^ currents, respectively. 1 μM tetrodotoxin (TTX) was used to block TTX-sensitive Nav channels.

Whole-cell voltage-clamp recordings were performed using a MultiClamp amplifier (Molecular Devices, 700B). Currents were filtered at 4 kHz (low-pass) and digitized at 20 kHz via a Digidata interface (Molecular Devices, 1440A). Series resistance was compensated by 80% to minimize voltage errors. Leak currents were subtracted by using an online P/4 procedure. The holding potential was set to -70 mV, approximating the resting membrane potential of cortical neurons. To assess sodium current density, a hyperpolarization prepulse to -110 mV (200 ms) was used to remove inactivation, followed by a 20-ms depolarization to 0 mV. Sodium current density (nA/pF) was calculated by normalizing peak current to bleb capacitance. To evaluate Nav channel slow inactivation, a single 5-s depolarizing pulse (0 mV) was applied to induce slow inactivation followed by test pulses (0 mV, 20 ms) separated by 10-ms recovery intervals at -70 mV. The fraction of available Nav channels was calculated by normalizing the peak current amplitude of each test pulse to the peak current measured at the onset of the 5-s depolarization.

### Co-immunoprecipitation in transfected HEK293T cells

In co-immunoprecipitation experiments, one protein-of-interest was tagged with an HA epitope and the other with a Flag epitope. The pCAGIG vector bearing either a HA or Flag tag alone was used as a control in paired constructs. HEK293T cells transfected with a pair of constructs were harvested 24 hours post-transfection for protein expression. Cells were lysed in ice-cold lysis buffer (20 mM Tris-HCl, 150 mM NaCl, 0.5% n-Dodecyl β-D-maltoside (DDM, Anatrace, D310), and protease inhibitor cocktail (Selleckchem, B14002)) with gentle agitation at 4 °C for 2 hours. Lysates were centrifuged at 20,000 x g for 20 minutes at 4 °C to remove insoluble debris. The supernatant was transferred to a fresh tube and kept on ice.

Protein concentration of supernatant was determined using a colorimetric bicinchoninic acid (BCA) assay (Beyotime, P0012S). For each immunoprecipitation (IP), 400 μg of proteins was incubated with 30 μL of anti-HA nanobody magnetic beads (AlfaLifeBio, KTSM1335) or anti-Flag magnetic beads (Bimake, B26102) at 4 °C with rotation for 1 hour. Beads were then separated with DynaMag spin and washed three times with wash buffer (20 mM Tris-HCl, 150 mM NaCl, 0.05% DDM). The captured proteins were eluted with 25 μL SDS-PAGE loading buffer containing 100 mM DTT, followed by heating at 45 °C for 10 min. Samples were stored at -20 °C prior to western blot analysis.

### Co-immunoprecipitation in mouse brain tissues

Brain tissues (∼100 mg per sample) were freshly dissected from wild-type or *Prrt2*-*V5* knockin mice, and homogenized on ice using glass homogenizers in 1 mL of Tris-buffered saline (TBS, 20 mM Tris-HCl, 150 mM NaCl, protease inhibitor cocktail). The homogenates were centrifuged at 800 x g for 10 min at 4 °C to remove nuclei and cell debris. Supernatants were subjected to centrifugation at 40,000 x g for 45 minutes at 4 °C to isolate membrane fractions. The resulting membrane-enriched pellets were resuspended in TBS and solubilized in lysis buffer containing 0.3% DDM for 3 hours at 4 °C. Lysates were then cleared by centrifugation at 20,000 x g for 20 minutes at 4 °C. Protein concentration was measured via BCA assay. For each IP, ∼200 μg of proteins was incubated with 30 μL of V5-Trap magnetic beads (Chromotek, v5tma-20) at 4 °C with rotation for 1.5 hours. Beads were washed three times with wash buffer (20 mM Tris-HCl, 150 mM NaCl, 0.15% DDM), and bound proteins were eluted with 25 μL SDS-PAGE loading buffer containing 100 mM DTT, followed by heating at 45 °C for 10 min. Eluates were kept at -20 °C until use in western blotting.

### Western blotting

Protein samples were separated using 4-12% PAGE Gel (Nanjing ACE Biotechnology, ET15412L) in MES-SDS running buffer at 140 V for 60 minutes. Proteins were transferred onto polyvinylidene difluoride (PVDF) membranes at 300 mA for 2 hours at 4 °C. The PVDF membranes were blocked in 5% nonfat milk in TBST (TBS with 0.1% Tween-20) for 1 hour at room temperature, followed by incubation with indicated primary antibodies overnight at 4 °C with gentle agitation.

After three washes in TBST, membranes were incubated with HRP-conjugated secondary antibodies for 2 hours at room temperature. Following another three washes, chemiluminescent substrate (TIANGEN Biotech, PA112) was applied, and signals were detected using a blot imaging system (Analytikjena, ChemStudio 815). Densitometric analysis of protein bands was performed using QuantityOne software (Biorad, Quantity One 1-D analysis, v4.6.2). The primary antibodies we used in western blotting were as follows: Anti-human PRRT2 (Atlas antibodies, HPA014447), Anti-Nav1.2 (Alomone Labs, ASC-002), Anti-Nav1.1 (Alomone Labs, ASC-001), Anti-HA-Tag (Cell signaling technology, 3724S), Anti-GAPDH-HRP (KangChen Bio-tech,KC-5G5), Anti-ATP1B2 (Abcam, ab185210), Anti-mouse PRRT2 (Wiiget Biotech, Rp3246) and Anti-β Actin-HRP (Cell signaling technology, 5125S).

### Animal Surgery

Mice were anesthetized with isoflurane (RWD life science, R510-22-10), using 4% for induction and 2% for maintenance. Body temperature was maintained throughout the procedure with a heating pad. Meloxicam (5 mg/kg, subcutaneous) was administered preoperatively for analgesia.

After securing the mice in a stereotaxic frame (RWD life science, Model 68528), 0.25% bupivacaine was locally injected at the incision sites for perioperative analgesia, and ophthalmic ointment was applied to protect the cornea. A midline incision was made to expose the skull, which was cleaned with 2% hydrogen peroxide using sterile cotton swabs to remove connective tissue. The skull surface was leveled to a stereotaxic plane, and the bregma was identified and marked. Once dry, the exposed skull was coated with a light-curing self-etch adhesive (3M ESPE Single Bond Universal).

For cerebellar electrical stimulation experiments, two small holes separated by 1.5 mm were drilled in the skull above the fourth/fifth cerebellar lobules [anterior-posterior (AP), -6.25 mm; medial-lateral (ML), ±0.75 mm; dorsal-ventral (DV), -1.0 mm from the dura]. Two metal electrodes (100 μm in diameter; California Fine Wire Company, M426550) were implanted and secured with dental cement.

For cortical electrical stimulation and EEG recording, two small craniotomies (∼0.6 mm in diameter) were made over the left sensory cortex (AP -0.8 mm; ML 1.5 mm) and visual cortex (AP -3.5 mm, ML 1.5 mm) for the insertion of bipolar stimulation electrodes (180 μm in diameter), which were implanted to a depth of -0.6 mm from the dura. Stainless steel screws were used as EEG electrodes: one was implanted into the right visual cortex (AP -2.25 mm, ML -1.5 mm), and the second into the right cerebellar vermis to serve as the reference. An additional screw was placed in the left cerebellar hemisphere as the ground. The electrodes and a custom-made head plate were affixed to the skull using dental cement.

Post-operatively, meloxicam (5 mg/kg, subcutaneously) was administered once daily for 3 consecutive days. Mice were allowed to recover for one week prior to electrical stimulation

### Cerebellar electrical stimulation and behavior assessment

Before cerebellar electrical stimulation, mice were habituated to the head-fixed platform for 30 minutes per day over three consecutive days. Electrical stimulation (300 μA, 60 Hz, 1-ms pulse width, 2 s duration) was delivered to the cerebellar cortex through a stimulus isolator (AMPI, ISO Flex) driven by a Master-8 stimulator. Mouse behavior was recorded with a video camera at a frame rate of 30 frames per second for 10 minutes.

Dystonia in mice were defined by the presence of one or more of the following abnormal postures or movements: prolonged forelimb elevation; persistent backward extension of the ears, tightly apposed to the head; an elongated trunk posture with the abdomen close to the ground and prolonged elevation or excessive extension of hindlimbs.

### Electrostimulation and EEG recording in the cerebral cortex

One week after surgery, mice were habituated to a custom-built head plate holder for 30 minutes per day over three consecutive days prior to testing.

For stimulation experiments, the pre-implanted electrodes were connected to a stimulus isolator (AMPI, ISO Flex). Electrical pulses (1 ms pulse width, 60 Hz frequency, 2 s duration) were delivered through the isolator and controlled via TTL signals from a pulse generator (RWD life science, R820). EEG signals (bandpass: 1-300 Hz) were amplified 1000-fold using an AC/DC differential amplifier (A-M Systems, Model 3000), digitized at 1 kHz with a Digidata (Molecular Devices, 1322A), and acquired using Clampex software (Molecular Devices, v10.7).

To assess cortical resilience in *Prrt2*-mutant versus wild-type mice, a stimulus-response protocol was employed using progressively increasing current intensities (0, 20, 40, 80, 120, 160, 200, 240, 280, 320 µA). Each stimulation was separated by a 24-hour interval. The stimulation threshold was defined as the lowest current intensity that induced after-discharges (lasting ≥ 2 seconds) in the EEG trace. Each EEG recording lasted for 5 minutes, with electrical stimulation delivered immediately following a 1-minute baseline period.

### Statistics

All grouped data are presented as mean ± standard error of the mean (s.e.m) unless otherwise stated. Statistical analyses were performed using Graphpad Prism 8 (GraphPad Software Inc, La Jolla, US). No statistical methods were used to pre-determine the sample sizes. The D’Agostino-Pearson omnibus (K2) test was used to assess normality.

Two-tailed, unpaired Student’s *t*-tests were applied in Figures 7B and 7D. A two-tailed, paired Student’s *t*-test was used in Figures 6D-F and Figure 7-figure supplement 1B. Two-tailed, unpaired Mann-Whitney test was applied in Figure 7H. Two-way ANOVAs were conducted for comparisons in Figures 1C-D, 1F-G, 2B-C, 2E-F, 3C-D, 3F-G, 4B-E, 4G-J, 7I, Figure 1-figure supplement 1B-C, Figure 3-figure supplement 1C-D, and Figure 7-figure supplement 2B-C. If necessary, the Geisser-Greenhouse correction was applied to adjust for violations of sphericity.

Statistical tests and sample sizes (including number of cells, slices, or animals) are stated in the figure legends. No data were excluded from analyses. Asterisks in figures denote significance levels: *P < 0.05, **P < 0.01, ***P < 0.001, ****P < 0.0001, n.s., not significant.

Co-immunoprecipitation assays were independently repeated at least three times in both *in vitro* and *in vivo* tests. All attempts at replicating experiments presented in the manuscript have obtained consistent conclusions.

Electrophysiological recordings were conducted with the experimenter blinded to group identity (transfected condition or animal genotype).

## Acknowledgements

The authors thank Fei Dong, Ting-Bin Ma, Xi Zhou, Wen Zhang, Wei Ke, Yue-Jie Xiao, Xue-Qin Jin and You-Sheng Shu for technical support for electrophysiological recording. We thank Huai-Zong Shen, Zhang-Qiang Li, Xue-Jing Huang and Shu-Jia Zhu for experimental support for protein expression. We thank Tong-Zhou Li, Hong Zhang, Jun-Jun Liu and Hai-Yan Zhen from ICE Bioscience for experimental support in sodium current recording. We appreciate the staff at the Core Facilities for their generous technical support, and members of Xiong Lab for technical support and insightful discussions.

This work was supported by Innovation of Science and Technology 2030-Major Project “platform of nonhuman primate models”, grant 2021ZD0200900 (Z.-Q.X.); National Natural Science Foundation of China, grants 82271269 (B.L.) and 82021001 (Z.-Q.X.); and Shanghai Municipal Science and Technology Major Project (Z.-Q.X.).

## Additional information

## Materials availability

All plasmids and mice generated in this study are available from the lead contact with a completed materials transfer agreement.

## Data and code availability

The data supporting the findings of this study are included in the main text. This paper does not report the original code.

## Contributions

Z.-Q.X. and B.L. conceived the study and designed the experiments. B.L., Q.-W.X. and J.Z. carried out cellular and molecular experiments and data analyses. X.-M.W., J.-Y.H. and J.-Q.P. performed animal experiments and data analyses. L.Z., J.Z., K.-X.L. and Z.-Y.W. contributed to plasmid construction. G.Y. was responsible for the knock-in mice generation. Y.-X.Z. helped with animal surgery and behavior analyses. B.L. wrote the manuscript. Q.-W.X., J.Z., X.-M.W., J.-Y.H. and Z.-Y.W. carried out manuscript review and editing. Z.-Q.X. and B.L. carried out project supervision, data interpretation and funding acquisition.

## Supplemental information

Figure 1-figure supplement 1, Figure 2-figure supplement 1, Figure 3-figure supplement 1, Figure 6-figure supplement 1, Figure 7-figure supplement 1 and Figure 7-figure supplement 2.

---

**Figure 1-figure supplement 1.**
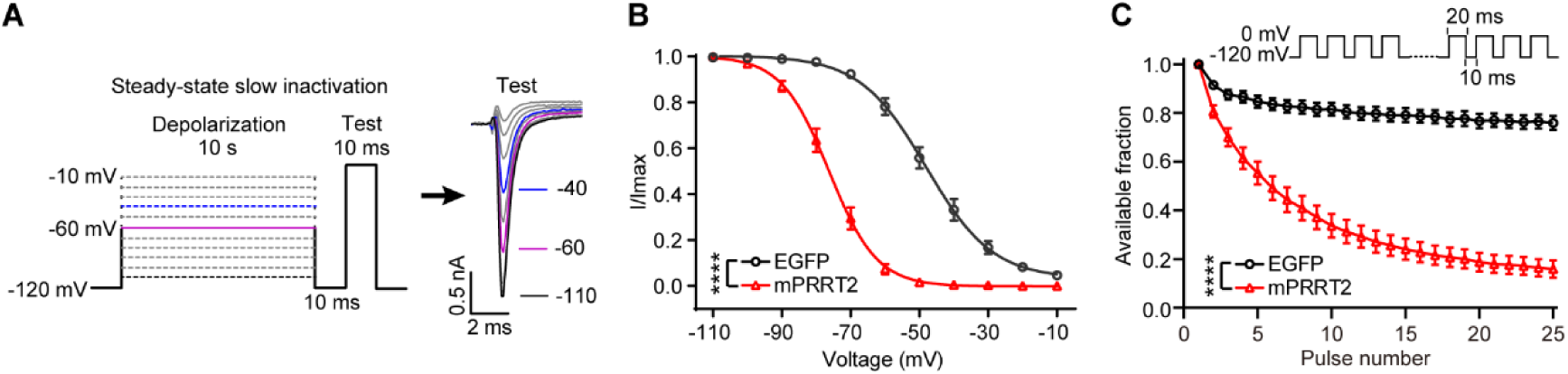
PRRT2 promotes steady-state slow inactivation and use-dependent inactivation. **(A)** Protocol of steady-state slow inactivation test (left) and representative traces of the sodium currents evoked by test pulses (right). The sodium currents gradually decreased as the voltage in conditioning steps increased. Scale bar, 0.5 nA, 2 ms. **(B)** The effects of PRRT2 on voltage-dependent steady-state slow inactivation of Nav1.2 channels (EGFP: n = 11 cells, mPRRT2: n = 13 cells). mPRRT2, mouse PRRT2. **(C)** PRRT2 modulates use-dependent inactivation of Nav1.2 channels during pulse trains (EGFP: n = 6 cells, mPRRT2: n = 6 cells). Data are presented as mean ± s.e.m. Two-way ANOVAs were used in (B and C) to determine the statistical significance of main effects of group. ****P < 0.0001.

**Figure 2-figure supplement 1.**
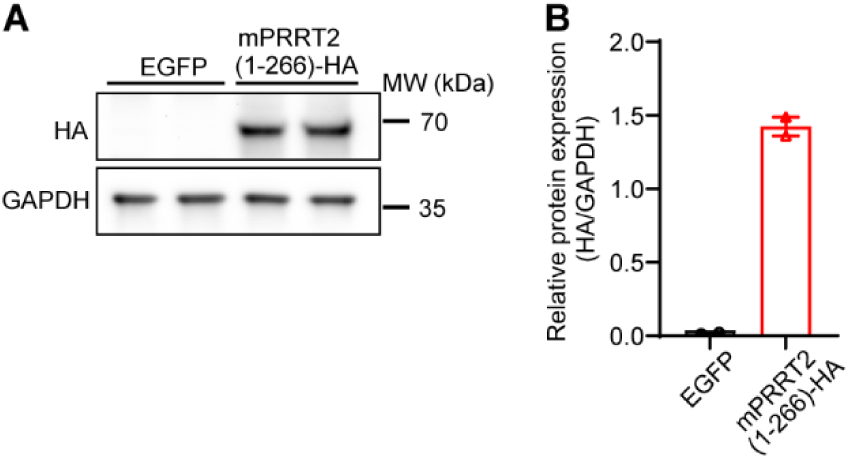
Western blot verification of mPRRT2(1-266) expression. **(A)** Representative Immunoblot showing HA-tagged mPRRT2 (1-266) expression in Nav1.2-stably expressing HEK293T cells. GAPDH served as a loading control. **(B)** Quantification of mPRRT2(1-266)-HA expression normalized to GAPDH (n = 2 independent samples). Data are presented as mean ± s.e.m.

**Figure 3-figure supplement 1.**
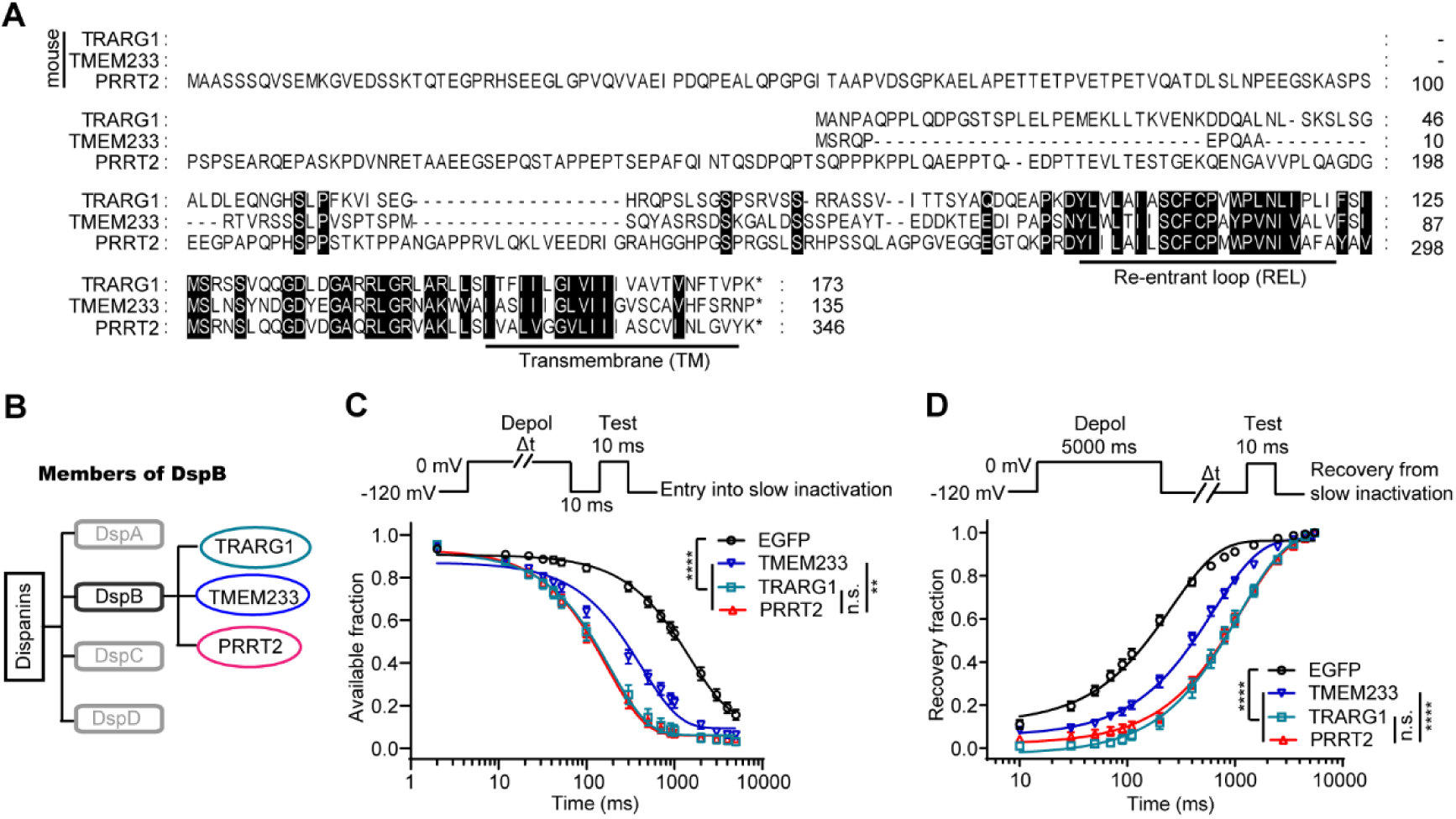
Paralogs of PRRT2 regulate Nav channel slow inactivation. **(A)** Sequence alignment of mouse TRARG1, TMEM233 and PRRT2 proteins. Conserved amino acids are highlighted and the transmembrane domains in carboxyl terminus of these proteins are underlined. REL, re-entrant loop. TM, transmembrane domain. **(B)** Schematic showing the phylogenetic relationship between members of Dispanin subfamily B (DspB). **(C)** The effects of members of DspB on the entry of Nav1.2 channels into slow inactivation (EGFP: n = 13 cells, TRARG1: n = 17 cells, TMEM233: n = 13 cells, PRRT2: n = 13 cells). **(D)** The effects of members of DspB on the recovery of Nav1.2 channels from slow-inactivated states induced by 5-s depolarization (EGFP: n = 10 cells, TRARG1: n = 14 cells, TMEM233: n = 12 cells, PRRT2: n = 12 cells). Data are presented as mean ± s.e.m. The main effect of group was assessed using two-way ANOVAs (C and D). **P < 0.01, ****P < 0.0001, n.s., not significant.

**Figure 6-figure supplement 1.**
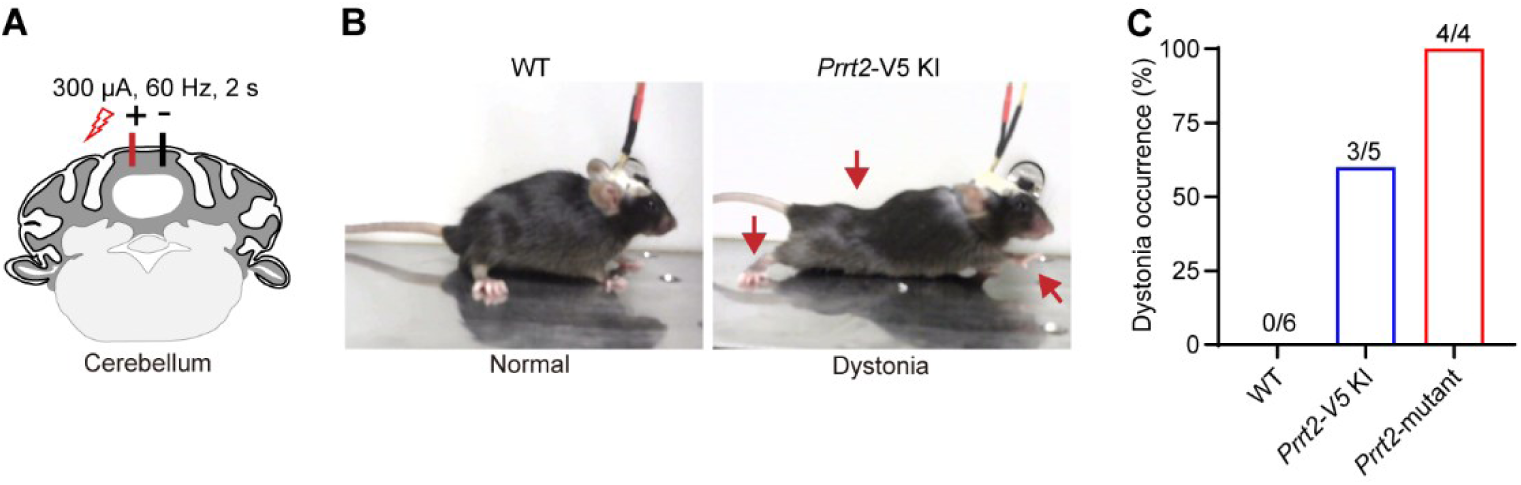
Dystonia behaviors in a subset of Prrt2-V5 knock-in mice. **(A)** Schematic illustration of electrical stimulation delivered to the localized area of the fourth/fifth cerebellar lobule in mice. **(B)** Representative postures of wild-type (WT) and Prrt2-V5 knock-in (KI) mice following cerebellar stimulation. Arrows indicate the body parts affected during the dystonic episode. **(C)** Incidence of dystonia behaviors induced by cerebellar electrical stimulation in WT, Prrt2-V5 KI, and Prrt2-mutant mice (WT: n = 6 mice, Prrt2-V5 KI: n = 5 mice, Prrt2-mutant: n = 4 mice). Data are presented as percentage of animals that exhibited dystonia behaviors within 5 minutes after stimulation.

**Figure 7-figure supplement 1.**
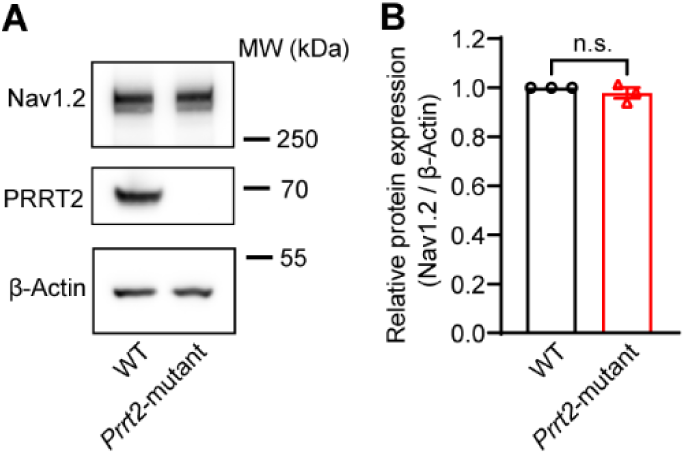
Protein levels of Nav1.2 are unchanged in brain tissue from Prrt2-mutant mice. **(A and B)** Immunoblots (A) and quantifications (B) of Nav1.2 proteins in brain tissue of wild-type (WT) and Prrt2-mutant mice (n = 3 mice). Data are presented as mean ± s.e.m. Two-tailed, paired Student’s t-test was used in (B) to determine the statistical significance. n.s., not significant.

**Figure 7-figure supplement 2.**
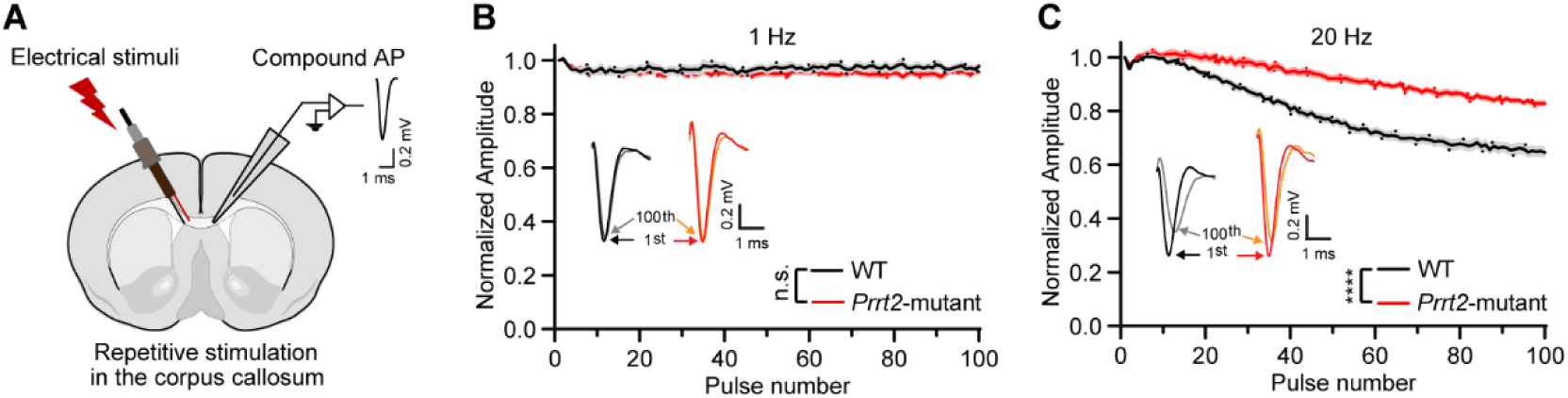
Activity-dependent adaptation is impaired in Prrt2-mu-tant mice. **(A)** Schematic illustration of corpus callosum stimulation and compound action potential (AP) recording in acute brain slices. Insert: representative trace of a compound AP recorded from the corpus callosum. Scale bar, 0.2 mV, 1 ms. **(B and C)** Normalized peak amplitude of compound APs in corpus callosum during repetitive electrical stimulation (100 μA) at 1 Hz (B) and 20 Hz (C) in WT and Prrt2-mutant mice (WT: n = 22 slices from 5 mice, Prrt2-mutant: n = 20 slices from 4 mice). Inserts: representative traces of the 1st and 100th compound APs recorded from the corpus callosum of WT and Prrt2-mutant mice. Scale bar, 0.2 mV, 1 ms. Data are presented as mean ± s.e.m. Two-way ANOVAs were used in (B) and (C) to determine the statistical significance of main effect of genotype. ****P < 0.0001. n.s., not significant.

